# Autism-associated transcriptional regulators target shared loci proximal to brain-expressed genes

**DOI:** 10.1101/2022.10.17.512583

**Authors:** Siavash Fazel Darbandi, Joon-Yong An, Kenneth Lim, Nicholas F. Page, Lindsay Liang, Athena R. Ypsilanti, Eirene Markenscoff-Papadimitriou, Matthew W. State, Alex S. Nord, Stephan J. Sanders, John L. R. Rubenstein

## Abstract

Many autism spectrum disorder (ASD)-associated genes act as transcriptional regulators (TRs). ChIP-seq was used to identify the regulatory targets of ARID1B, BCL11A, FOXP1, TBR1, and TCF7L2, ASD-associated TRs in the developing human and mouse cortex. These TRs shared substantial overlap in the binding sites, especially within open chromatin. The overlap within a promoter region, 1-2,000bp upstream of transcription start site, was highly predictive of brain expressed genes. This signature was observed at 96 out of 102 ASD-associated genes. In vitro CRISPRi against ARID1B and TBR1 delineated downstream convergent biology in mouse cortical cultures. After eight days, NeuN+ and CALB+ cells were decreased, GFAP+ cells were increased, and transcriptomic signatures correlated with the postmortem brain samples from individuals with ASD. We suggest functional convergence across five ASD-associated TRs leads to shared neurodevelopmental outcomes of haploinsufficient disruption.

## Introduction

Autism spectrum disorder (ASD) is a common and highly heritable neurodevelopmental disorder (*1*). To date, over a hundred genes have been associated with ASD, mostly through the detection of rare loss-of-function variants that disrupt the function of one of the two copies of a gene (*2, 3*). However, the mechanism by which disruption of these genes leads to ASD symptoms remains elusive. Analysis of patterns of gene expression for these ASD-associated genes in the developing human brain has implicated excitatory and inhibitory cortical and striatal neurons (*2–6*). Orthogonal analysis of the *postmortem* brain in ASD cases identifies down-regulated gene expression modules, which are enriched for both neuronal marker genes and ASD-associated genes, and up-regulated gene expression modules enriched for non-neuronal marker genes but not ASD-associated genes (*7–9*). Mouse experiments observed similar transcriptomic patterns as a consequence of disrupting multiple ASD-associated genes, with some gene expression profiles overlapping those seen in the *postmortem* ASD brain (*10*). These results suggest that convergent pathology, captured by high-dimensional transcriptomic datasets, may underlie the shared phenotypic consequences across multiple ASD-associated genes. This, in turn, raises the question of how disrupting multiple genes with heterogeneous and pleiotropic functions can yield similar transcriptomic and phenotypic outcomes.

The majority of ASD-associated genes encode proteins that act as transcriptional regulators (TRs), influencing the expression of other genes; these include transcription factors (e.g., *TBR1*, *FOXP1*), histone modifiers (e.g., *KMT5B*), and chromatin remodelers (e.g., *CHD8*) (*2*). The genomic targets of these regulatory genes and the transcriptomic consequences of their disruption remain largely uncharacterized, as do their functional relationship to other ASD-associated genes. Identifying these genomic targets could reveal convergent gene regulatory networks and predict downstream neurobiology to account for the shared autistic phenotype; they could also provide an orthogonal approach to distinguish the cell types, brain regions, and developmental stages involved. Analyses of the genomic targets of individual genes support this possibility, for example, targets of CHD8, POGZ, and TBR1 are enriched for ASD-associated genes (*11–14*), however, identification of shared targets is complicated by heterogeneous protocols, species, cells/tissues, and developmental stages.

To assess the extent of shared regulatory targets across ASD-associated genes, we selected five TRs for further analysis, based on strong evidence for ASD association (Fig. 1A), expression during cortical development, evidence of direct binding to DNA, and the availability of reliable antibodies*. ARID1B* and *BCL11A* are both DNA-binding subunits of the BAF (SWI/SWF) chromatin remodeling complex expressed across multiple tissues (*15, 16*), though cortical *BCL11A* expression appears restricted to neurons (*17, 18*). Both *FOXP1,* a winged-helix TR, and *TBR1,* a T-Box TR, are expressed highly in neurons of the developing cortex, *FOXP1* is also expressed in neurons of the developing striatum (*17–20*). *TCF7L2* is an HMG-Box TR expressed in the cortical and subcortical progenitors (*21*) and thalamic excitatory neurons (*22*). We observed substantial overlap in the genomic targets of all five TRs in the developing human and mouse cortex, especially in proximity to genes highly expressed in the cortex, including most ASD-associated genes. These shared genomic targets are proximal to genes critical to brain development and function, and thus suggest a mechanism for similar phenotypic consequences for mutations in the diverse set of ASD genes.

**Figure 1.**
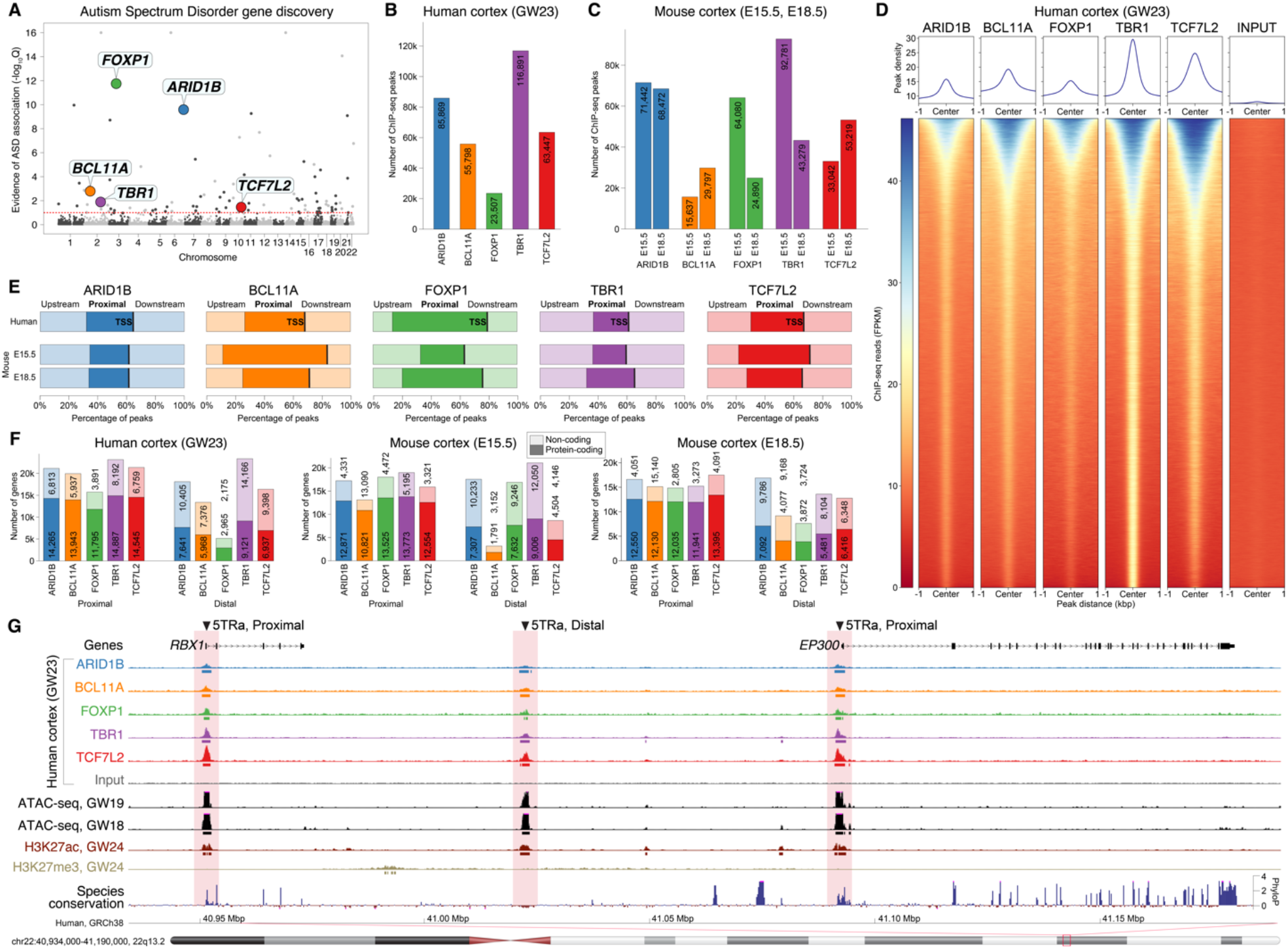
ChIP-seq peaks identified in five ASD-associated TRs. **A)** Evidence of ASD association for five DNA-binding TRs from exome sequencing (*2*). **B)** ChIP-seq peaks were identified in human cortex at gestation week 23 (GW23) and **C)** mouse cortex (embryonic days 15.5 and 18.5). **D)** Read counts around ChIP-seq peaks for human cortex. The union of peaks across the five TRs is shown in the same order (y-axis) for all six datasets (additional datasets shown in Fig. S1). **E)** ChIP-seq peaks for all five transcription factors are enriched for promoter regions ≤2,000bp proximal to the transcription start site (TSS). **F)** ChIP-seq peaks are found proximal to the TSS of 10,663 to 14,874 protein-coding genes across species and assay. **G)** A representative example of TR peaks in human prefrontal cortex proximal and distal to two genes are shown alongside peaks from ATAC-seq in GW18 and GW19 human prefrontal cortex and histone ChIP-seq data (H3K27ac, H3K27me3) in GW24 human prefrontal cortex (compared to an ATAC-seq only region, Fig. S5). Abbreviations: 5TRa: locus with binding by all five ASD-associated TRs (5TR) and has an ATAC-seq peak (a); E15.5/E18.5: embryonic day 15.5/18.5; GW: gestation week.

### Identifying regulatory targets of five ASD-associated TRs in developing human and mouse neocortex

We generated chromatin immunoprecipitation sequencing (ChIP-seq) data for these five ASD-associated TRs from human frontal neocortex at gestational week 23 (GW23) and mouse neocortex at embryonic day (E)15.5 and E18.5 (Table S1). Peak counts ranged from 23,507 to 116,891 (Fig. 1B-C), were distinct from reads in input and blocking peptide controls (Figs. 1D, S1), were consistent across biological replicates and with previously published data (*13*) (Fig. S2), and showed substantial conservation across species (Fig. S3). Four of the TRs have known motifs (BCL11A, FOXP1, TBR1, and TCF7L2), which were enriched in the corresponding ChIP-seq peaks compared to scrambled sequence (Fig. S4). In contrast, ATAC-seq peaks were present at numerous loci without TR peaks (Figs. S1, S5). A high proportion of ChIP-seq loci were proximal peaks, defined as overlapping the promoter region, mapping 0-2,000 bp upstream of any transcription start site (TSS), compared to distal peaks, defined as peaks not overlapping promoters (Fig. 1E). Despite considerable variation in peak counts (Fig. 1B-C), the number of protein-coding genes with a proximal peak was similar between TRs, species, and developmental stages, ranging from 10,663 to 14,874 genes (Fig. 1F).

### ASD-associated TRs converge on a common set of targets

The ChIP-seq peaks for the five ASD-associated TRs frequently targeted the same genomic loci in developing human cortex (Fig. 1D, 1G), suggesting shared regulatory networks and/or protein complexes (*23*). To assess the expected degree of overlap between TRs in heterogeneous tissues, we reprocessed ENCODE ChIP-seq data for 14 TRs in the adult human liver through our analysis pipeline (Table S2). We used p-values to identify the top 10,000 proximal peaks for each TR and assessed the intersection and correlation by p-value rank for all combinations of TRs. The ASD-associated genes showed a greater degree of both correlation (0.42 Spearman’s rho in cortex vs. 0.07 in liver, P = 1.3 x 10^-6^, Wilcoxon; Fig. 2A) and intersection (83.8% percent intersecting peaks in cortex vs. 59.2% in liver, P = 2.7 x 10^-7^, Wilcoxon; Fig. S6A) than TRs in the liver, except for CTCF and RAD21, (components of the chromatin looping complex (*24*)). Performing this analysis for 8,000 distal peaks also showed high overlap, especially between ARID1B, BCL11A, and TBR1 (0.26 Spearman’s rho in cortex vs. 0.13 in liver P=0.04 for correlation, Fig. 2B; 46.1% percent intersecting peaks in cortex vs. 36.0% in liver, P=0.37, Fig. S6B).

**Figure 2.**
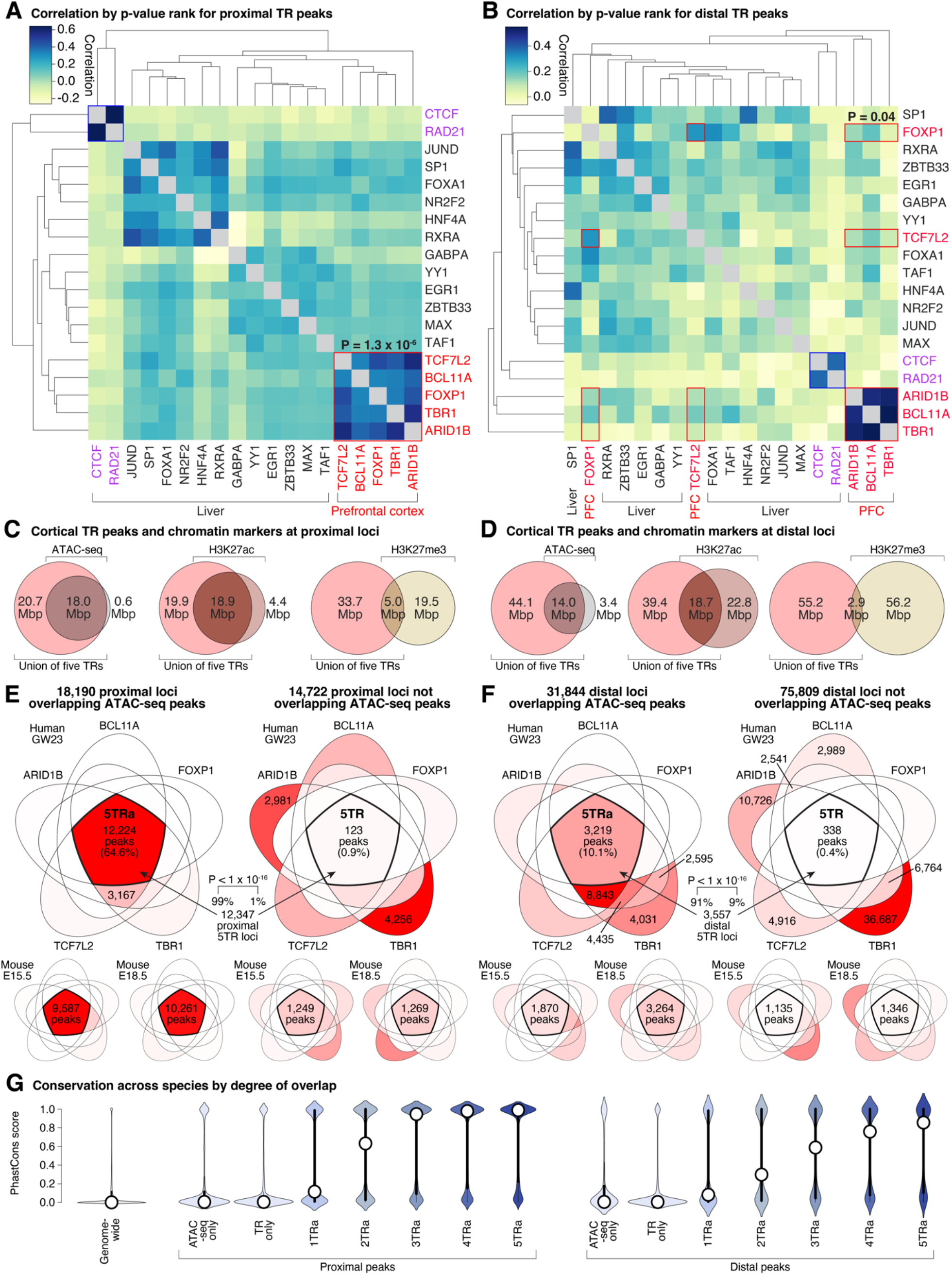
Overlap between ChIP-seq peaks from five ASD-associated TRs. **A)** Top 10,000 proximal ChIP-seq peaks ranked by p-value from 14 TRs in adult human liver (ENCODE, Table S2) and five ASD-associated TRs in fetal human cortex (Table S1) to assess intersection (Fig. S6A) and correlation of peak ranks. **B)** Equivalent plot for 8,000 distal peaks. **C)** Overlap between proximal ChIP-seq peaks for five ASD-associated TRs and three epigenetic markers in human fetal cortex. **D)** Equivalent overlaps for distal peaks. **E)** Overlap between proximal ChIP-seq peaks for the five ASD-associated TRs in developing human cortex overlapping (top left) or not overlapping (top right) with ATAC-seq peaks. Color gradient represents the percentage of peaks in each section, with red being the highest percentage and white being 0%; peak counts are given for the intersection of all five TRs and sections with greater than 1,500 peaks. Equivalent plots are shown for E15.5 mouse cortex and E18.5 mouse cortex (bottom), and **F)** distal peaks. **G)** PhastCons scores for conservation across 100 vertebrate species are shown genome-wide (left), for loci with ATAC-seq peaks but no ChIP-seq TR peaks or ChIP-seq TR peaks by no ATAC-seq peaks, and for ATAC-seq peaks intersecting with one to five ChIP-seq TR peaks (1TRa – 5TRa). Abbreviations: 5TRa: peak with all five ASD-associated transcription factors (5TR) and ATAC-seq (a); GW23: gestational week 23; E15.5/E18.5: embryonic day 15.5/18.5. Statistical analyses: A and B: Wilcoxon test; E: Permutation test.

To further understand the implications of these overlapping TR peaks in developing human cortex, we considered the overlap with open chromatin regions, detected by the assay for transposase-accessible chromatin sequencing (ATAC-seq) in GW18/19 human cortex (*25*), and H3K27ac and H3K27me3 histone modifications detected by ChIP-seq in GW24 human cortex (Fig. 2C-D). We observed substantial overlap between the proximal ASD-associated TR peaks and ATAC-seq (55.5% of TR peaks and 46.5% of nucleotides covered by TR peaks) and H3K27ac peaks (53.4% of peaks and 48.7% of nucleotides), both of which are associated with active transcription. In contrast, overlap with H3K27me3, a marker of gene repression, was minimal (16.5% of peaks and 13.0% of nucleotides). A similar pattern was observed in developing human cortex for distal peaks (Fig. 2D). Thus, the five ASD-associated TRs predominantly bind to proximal and distal loci with epigenetic states indicative of active transcription.

We next considered whether overlaps between ChIP-seq peaks of multiple ASD-associated TRs (Fig. 1G, 2A-B) occurred within the ATAC-seq-defined open chromatin regions or not (Fig. 2C-D, Fig. S7). In developing human cortex, we identified 32,962 independent proximal loci targeted by one or more ASD-associated TRs, split approximately in half between those with concurrent ATAC-seq peaks (18,190, 55.2%) and those without (14,772, 44.8%). Of the 32,962 proximal loci, 12,347 (37.5%) are targeted by all five ASD-associated TRs (5TR) and, remarkably, almost all of these have concurrent ATAC-seq peaks (12,224, 99.0% of 5TR proximal peaks, 64.6% of all proximal peaks; P<1×10^-10^, permutation testing to account for size differences, Fig. 2E). In contrast, of the 10,736 proximal loci targeted by a single ASD-associated TR, only 882 (8.2%) had concurrent ATAC-seq peaks. Similar enrichment for overlapping ASD-associated TR ChIP-seq peaks within ATAC-seq peaks were observed for mouse cortical data at both E15.5 and E18.5 (Fig. 2E) and for developing human cortex data within H3K27ac peaks (Fig. S6C-D).

Enrichment for overlapping ASD-associated TR ChIP-seq peaks was also observed within ATAC-seq peaks for distal peaks (Fig. 2F). The 107,653 independent distal peaks included 31,844 (29.6%) with concurrent ATAC-seq peaks and 75,809 (70.4%) without. Out of 3,557 distal loci targeted by all five ASD-associated TRs, 3,219 (90.5%) have concurrent ATAC-seq peaks (P<1×10^-10^, permutation testing, Fig. 2F), while most of the 62,474 distal loci targeted by a single ASD-associated TR occur outside of ATAC-seq-marked open chromatin regions (55,911, 89.5%). There were 6,063 distal loci identified by ATAC-seq without any of the five ASD-associated TRs. As with proximal loci, similar patterns are seen in mouse cortical data (Fig. 2F) and for H3K27ac peaks in developing human cortex (Fig. S6E-F).

### Open chromatin regions targeted by all five ASD-associated TRs are highly conserved

Given the high degree of overlap between ATAC-seq peaks and ChIP-seq peaks from all five ASD-associated TRs, hereafter referred to as “5TRa”, we sought to characterize these regions in depth. The 12,224 proximal 5TRa loci in the developing human cortex span 26.2Mbp and are upstream of the transcription start site of 11,695 protein-coding transcripts and 4,739 non-coding transcripts (Table S3). ChIP-seq peaks for each TR contributing to the 5TRa regions were called with higher confidence, based on lower p-values, than peaks outside the 5TRa regions (Fig. S6G). Most of these proximal loci are highly conserved across species, with 8,233 (67.4%) including a region with a max PhastCons score above 0.5 (Fig. 2G). Moreover, 8,867 (72.5%) overlap with a 5TRa peak in mouse E15.5 cortex, and 8,639 (70.7%) overlap with a 5TRa proximal loci in mouse E18.5 cortex.

The 3,219 distal 5TRa regions from developing human cortex span 5.7Mbp. Based on the nearest transcription start site, these loci are related to 850 protein-coding transcripts and 1,932 non-coding transcripts (Table S3). As seen for proximal loci, the distal 5TRa loci have lower p-values than those outside of 5TRa loci (Fig. S6H). Most are highly conserved across species, with 1,880 (58.4%) loci including a region with a max PhastCons score above 0.5 (Fig. 2G). Furthermore, 652 (20.3%)/548 (17.0%) overlap 5TRa distal loci in mouse E15.5/E18.5 cortex.

### ASD-associated TR-bound loci are enriched for motifs of TR genes associated with other neurodevelopmental and psychiatric disorders

Since the five ASD-associated TRs have DNA-binding domains, we used HOMER to assess whether DNA sequence motifs were enriched in the 5TRa proximal and distal loci against both genomic background and representative proximal/distal backgrounds (Fig. 3). Similar results were obtained using 5TRa loci from developing human and mouse cortex. Substantial enrichment was observed for motifs related to promoter-enhancer loops, including ZNF143/Staf and THAP11/Ronin in proximal elements and CTCF and CTCFL/Boris in distal elements (Fig. 3). Proximal 5TRa loci were also enriched for the ETS, KLF/SP, YY1, NKRF/Nrf, and NFY motif groups, while distal loci were enriched for the HTH/RFX and the bHLH motif groups. Many of the genes in these 5TRa-enriched motif groups are associated with neurodevelopmental and psychiatric disorders (10 out of 182 genes; 3.4-fold enrichment; *X*^2^ (1, N = 19,654) = 15.9, p=7×10^-5^; Table S4), including ASD (*2*): *RFX3* (HTH/RFX), *TCF4* (bHLH), *NCOA1* (bHLH), neurodevelopmental delay (*26*): *CTCF* (CTCF/BORIS), *YY1* (YY1), *ERF* (ETS), *KLF7* (KLF/SP), *TFE3* (bHLH), and *MYCN* (bHLH), and schizophrenia (*27*): *SP4* (KLF/SP).

**Figure 3.**
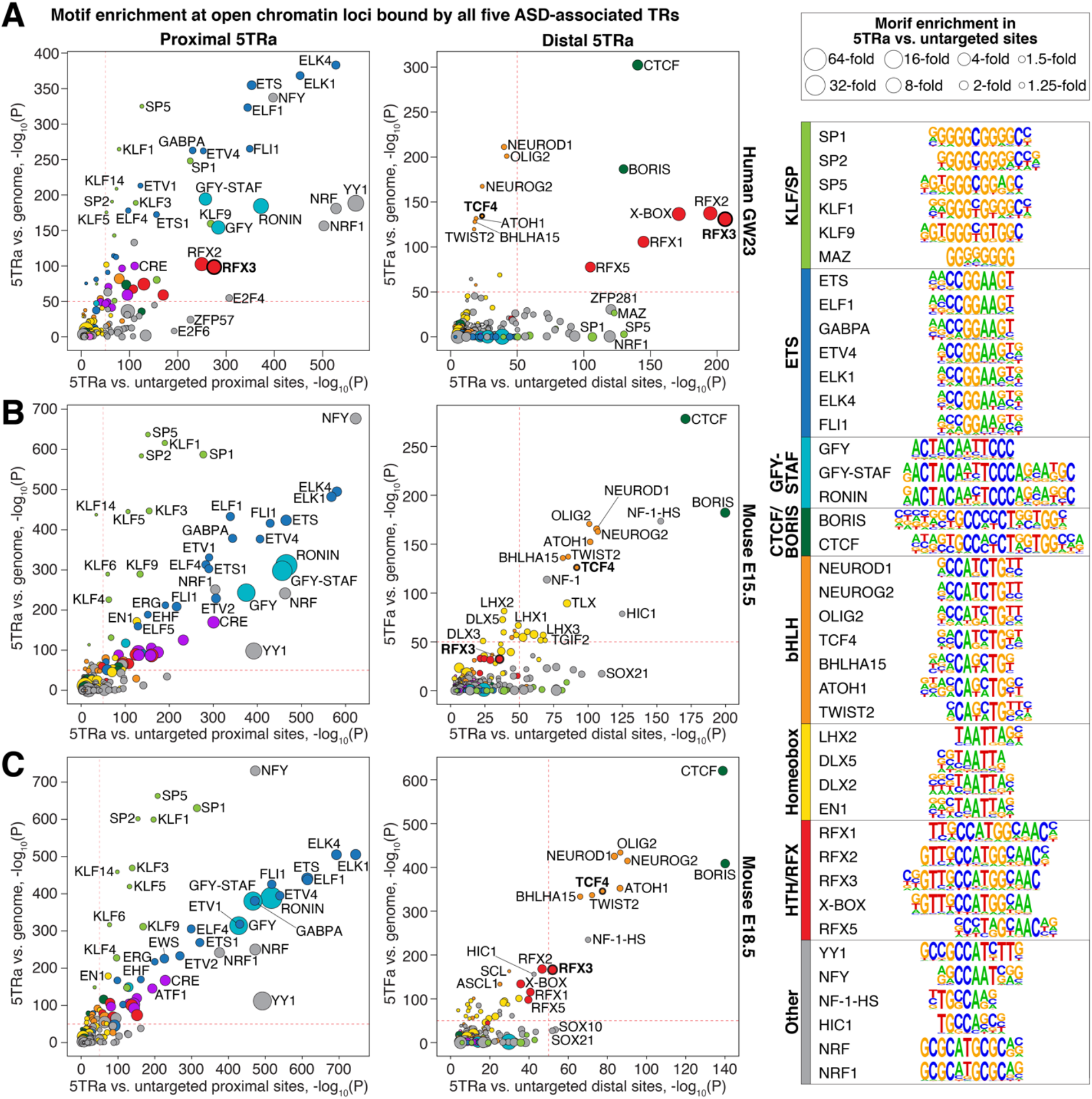
Motif enrichment in regions targeted by ASD-associated TRs. **A)** Results of HOMER known motif enrichment (Table S4) for proximal (left) and distal (right) loci bound by all five ASD-associated TRs and ATAC-seq (5TRa) against genomic background (y-axis) and against untargeted proximal/distal sites (x-axis, Supplementary Methods). Groups of related motifs are shown by color (panel on right) and motifs for ASD-associated genes (e.g., *RFX3*) are in bold font. Dashed red lines are included at P=1×10^-50^ to aid comparisons across panels. These analyses are repeated for mouse 5TRa loci at **B)** E15.5 and **C)** E18.5. Abbreviations: bHLH: Basic helix-loop-helix; CTCF: CCCTC-binding factor; ETS: E-twenty-six transformation-specific; GFY: General Factor Y; HIC1: HIC ZBTB transcriptional repressor 1; HTH: Helix-turn-helix; KLF: Krüppel-like family; NFY: Nuclear transcription factor Y; RFX: Regulatory Factor binding to the X-box; SP: Specificity protein; STAF: Selenocysteine TRNA Gene Transcription-Activating Factor (ZNF143). Statistical analyses: A-C: HOMER binomial enrichment.

### ASD-associated TRs are proximal to ASD-associated genes and brain-expressed genes

Prior analyses have described enrichment of ASD-associated genes in proximity to the binding sites of individual ASD-associated transcriptional regulators (*11–14*). We observed a similar result with proximal 5TRa loci from the developing human cortex upstream of 96 out of 102 ASD-associated genes (*28*) (94.1% of ASD genes vs. 65.7% of non-ASD genes; *X*^2^ (1, N = 17,484 all autosomal protein-coding genes) = 36.2, p=2×10^-9^; Fig. 4; Table S5). For the remaining six ASD-associated genes, five had overlapping peaks from at least three of the ASD-associated TRs proximal to their TSS. Equivalent results were also observed using data from mouse at E15.5 and E18.5 (Fig. 4F-G). Assessing ASD-enrichment by permutation test, accounting for gene cDNA length (a predictor of ASD gene discovery), yielded a similar result (P = 5×10^-4^).

**Figure 4.**
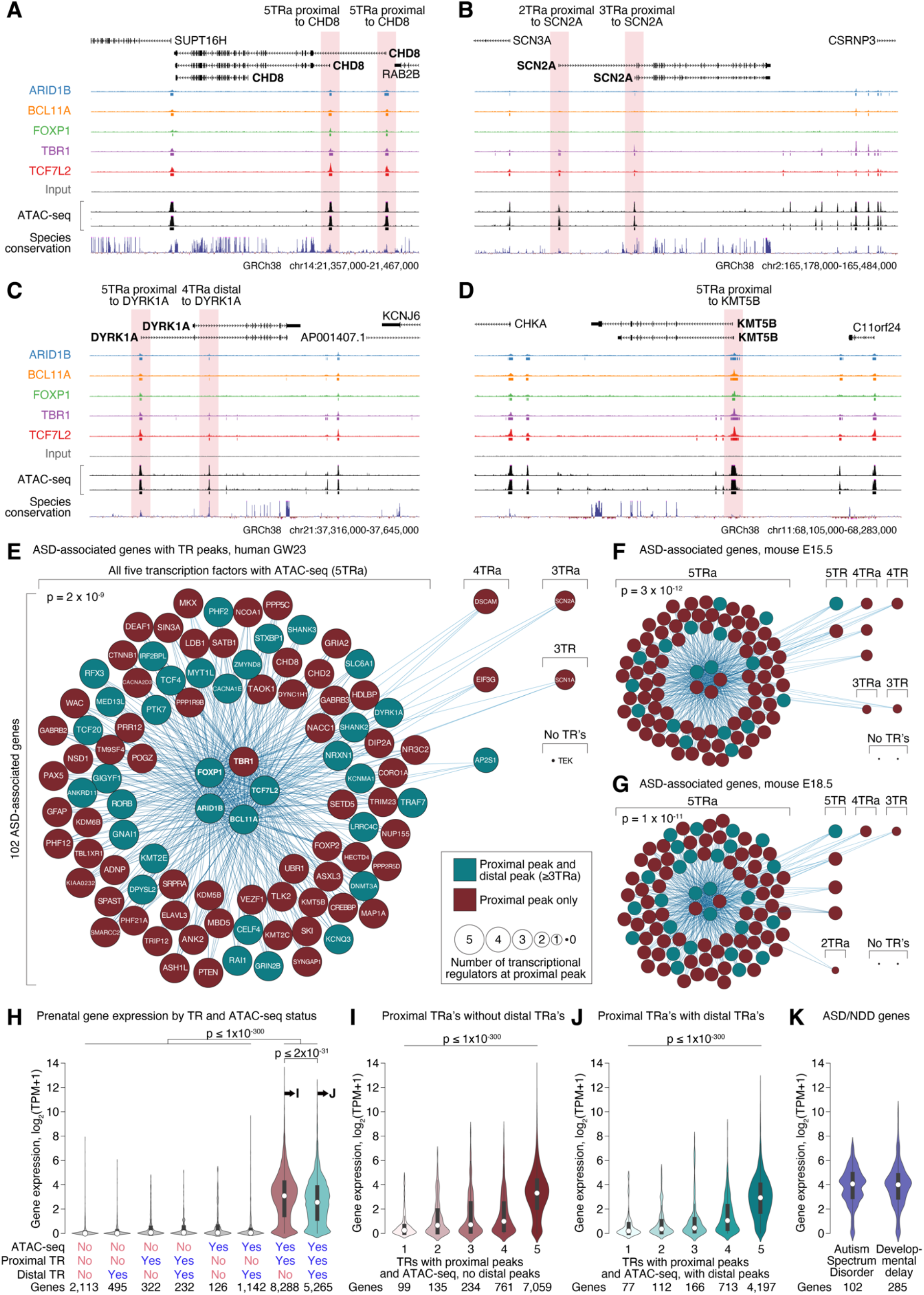
Enrichment of overlapping TR peaks at ASD-associated genes. A-D) TR peaks around the ASD-associated genes *CHD8*, *SCN2A*, *DYRK1A*, and *KMT5B*. **E)** Network plot showing whether ChIP-seq peaks for the five ASD-associated TRs (central circles/nodes) and ATAC-seq are detected proximal to the other 97 ASD-associated genes (peripheral circles/nodes, Table S5). Genes that also have a nearby distal peak that includes ARID1B, BCL11A, TBR1, and ATAC-seq (Fig. 2C, 2H) are shown in teal, while those without such a peak are in brown. Equivalent plots are shown for the same genes for the mouse data at **F)** E15.5 and **G)** E18.5. **H)** Median gene expression in the fetal human prefrontal cortex is represented for protein-coding genes binned by the presence or absence of at least one TR proximally or distally with or without ATAC-seq. **I)** Median gene expression in the fetal human prefrontal cortex is shown for all cortex-expressed protein-coding genes, binned by the number of ASD-associated TRs bound proximally in the absence of a distal TR and **J)** In the presence of a distal TR. **K)** Equivalent expression of genes associated with ASD or developmental delay. Abbreviations: 5TRa: peak with all five ASD-associated TRs and ATAC-seq, TPM: transcripts per million (a). Statistical analyses: E-G: Chi-squared; H-J: Logistic regression.

We next considered whether the presence of an ATAC-seq peak, or proximal ASD-associated TR peak, predicted gene expression levels in developing (GW12-40) human prefrontal cortex (*6*). Remarkably, most genes with proximal ATAC-seq peaks and at least one ASD-associated TR peak were robustly expressed (log_2_(TPM+1) ≥ 1), whereas almost all genes without both a proximal ATAC-seq and at least one ASD-associated TR peak were weakly expressed (log_2_(TPM+1) < 1, Fig. 4H). For peaks with both a proximal ATAC-seq peak and an ASD-associated TR peak, the number of ASD-associated TRs was highly predictive of the level of gene expression; this relationship was non-linear on a logarithmic scale of expression, with much higher expression in the presence of and ATAC-seq peak and all five ASD-associated TRs than four or fewer (Fig. 4I-J). Genes associated with ASD are highly brain expressed (median log_2_(TPM+1) = 4.0, Fig. 4K) and correcting for gene expression in the brain accounts for the extent of their enrichment in 5TRa loci, suggesting that the binding of multiple ASD-associated TRs proximal to ASD genes may contribute to their high brain expression.

### Genes with ASD-associated TR peaks have higher expression in the fetal brain

Target genes of distal loci were identified using a nearest TSS approach, including all protein-coding and non-coding transcripts. Given the overlap between ARID1B, BCL11A, and TBR1 at distal ATAC-seq peaks (Fig. 2), we assessed the number of ASD-associated genes with distal loci in developing human cortex containing at least these factors (≥3TRa). The 102 ASD-associated genes (*28*) were also enriched for these distal peaks (35.3% of ASD genes vs. 12.1% of non-ASD genes; *X*^2^ (1, N = 17,484 all autosomal protein-coding genes) = 50.5, p=1×10^-12^; Fig. 4; Table S5). However, the presence of a distal locus was associated with slightly lower gene expression in the developing human prefrontal cortex (Fig. 4H, 4J).

The enrichment we observed for GFY-STAF/ZNF143/THAP11 and CTCF/BORIS/CTFL motifs at 5TRa loci suggests that the 5TRa loci participate in chromosomal looping. Consistent with this idea, we identified potential interactions between ≥3TRa distal peaks and 77 of the 102 (75.5%) ASD-associated genes (Table S5) using the Activity-by-Contact (ABC) approach (*29*) with chromatin accessibility and Hi-C data from the developing human brain (*25, 30*). Similarly, a distal ≥3TRa peak was detected within 100kb of 70 ASD-associated genes (68.6%) and within 500kb for 96 (94.1%).

To better understand the function of these distal peaks, we considered the overlap with experimentally validated enhancer loci defined by VISTA (*31*). Of the 998 VISTA human elements with activity in E11.5 mice, 53 overlap with a 5TRa region, and 140 overlap with a ≥3TRa region that includes ARID1B, BCL11A, and TBR1 (Fig. 2B). As expected for a dataset derived from cortex, we observed enrichment for VISTA elements that are active in the telencephalon (63 regions, OR=2.17, p=5×10^-5^), including elements specific to both the pallium and subpallium in the E11.5 mouse (Fig. S8, Table S6). Two of these VISTA-positive regions also showed evidence of interaction with ASD genes through the ABC data: hs399 with *BCL11A* and hs416 with *ARID1B* (Table S6).

Next, we directly tested whether the hs399 VISTA region is regulated by TBR1. hs399 is ∼340kbp downstream of the *BCL11A* TSS and is bound by ARID1B, BCL11A, TBR1, and TCF7L2 in an open chromatin region (4TRa) (Fig. 5A). ABC analysis showed a potential interaction between this locus and the *BCL11A* TSS (Fig. 5A). We generated a stable hs399 enhancer transgenic mouse (hs399-CT2IG) that expresses GFP and Cre^ERT2^ (*32*). Immunohistochemistry shows that this regulatory element is active in the prenatal and postnatal developing cortex (Fig. 5B and not shown). However, in a constitutive *Tbr1^nul/null^* background hs399 enhancer activity is substantially reduced (Fig. 5B”). Furthermore, CRISPRi targeted to the hs399 element in cultured neonatal mouse cortical neurons resulted in an ∼10-fold decrease in *BCL11A* RNA expression (Fig. 5C).

**Figure 5.**
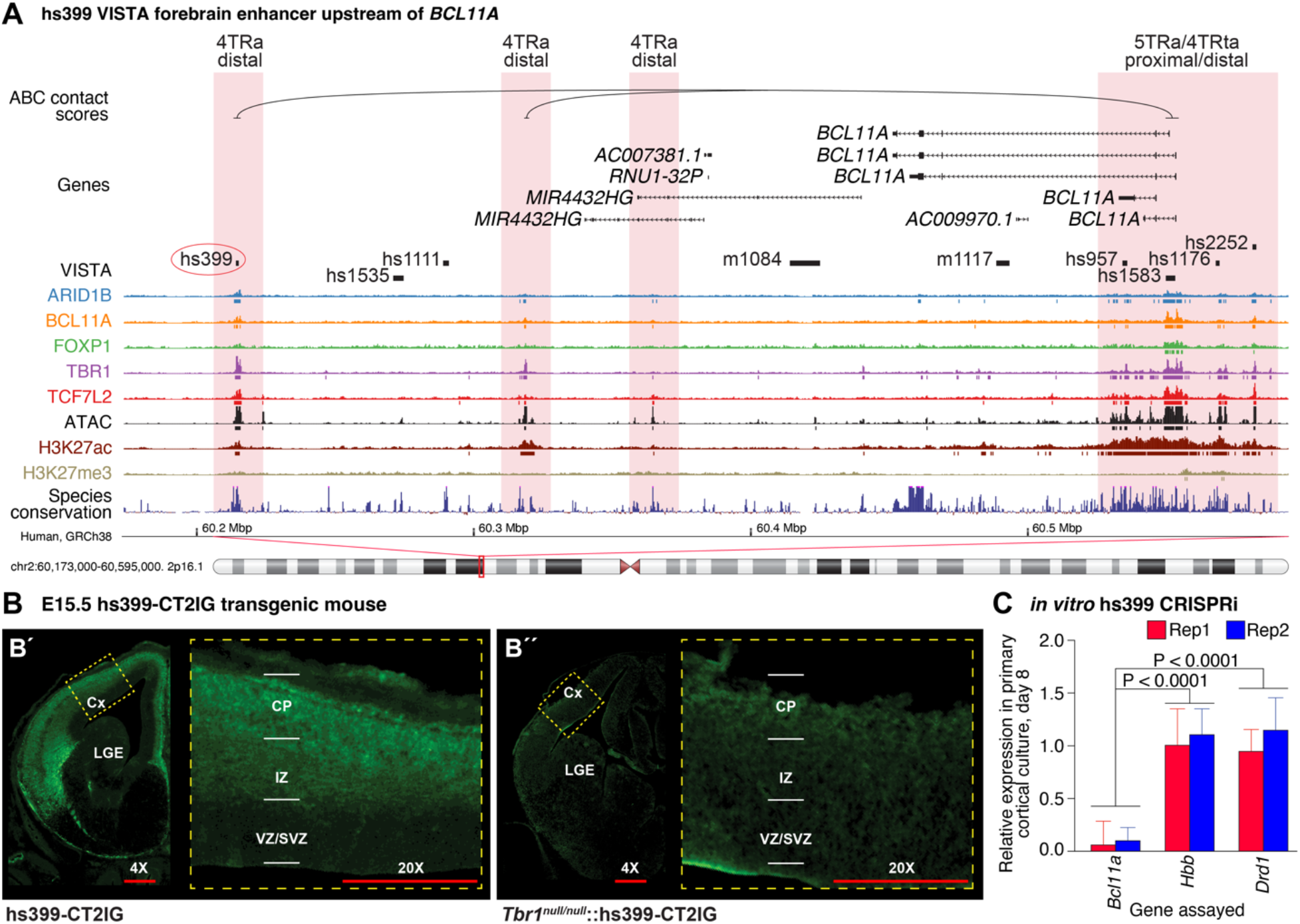
hs399 distal locus regulates *Bcl11a* expression in the developing mouse cortex. **A)** Four TRs encoded by ASD-associated genes (ARID1B, BCL11A, TBR1, and TCF7L2) bind to Vista element hs399 in the human prefrontal cortex at GW23, overlapping with ATAC-seq and H3K27ac ChIP-seq peaks. ABC data show a relationship with the *BCL11A* transcription start site 340,000bp downstream. **B)** hs399 is active in the cortical plate and intermediate zone of an hs399-CT2IG enhancer transgenic mouse at E13.5. TBR1 promotes the activity of the hs399 putative regulatory element, as the hs399 activity (GFP expression) is reduced in *Tbr1^null/null^*. Anti-GFP immunostaining is in green. **C)** qPCR analysis of *in vitro* CRISPRi guide RNA-directed against hs399 enhancer in mouse primary neocortical cultures, eight days post-transduction. CRISPRi directed to the hs399 locus decreased *Bcl11a* expression but did not impact *Hbb* and *Drd1* expression. Statistical analysis: Two-tailed T-Test with Tukey correction was used for pairwise comparisons. Abbreviations: Cx: cortex; CP: cortical plate; IZ: intermediate zone; LGE: lateral ganglionic eminence; SVZ: subventricular zone; VZ: ventricular zone. 4X and 20X refer to relative magnification.

### ASD-associated TRs target genes whose expression is enriched in cortical progenitors and neurons

ASD-associated genes are enriched within genes expressed by excitatory and inhibitory neurons from the human fetal cortex (*6, 17, 28*). Analysis of cells that express genes near 5TRa loci provides an orthogonal approach to identify the cell types involved in ASD. Therefore, we assessed cell type enrichment in ∼40,000 cells from the human fetal cortex at 17-18 weeks of gestation (*33*). Across 5TRa loci in human and mouse fetal cortex, enrichment was seen for cortical progenitor cells and excitatory neurons at proximal loci, and for excitatory and inhibitory neurons at distal loci (Fig. 6A). Consistent results were observed for other fetal cortex transcriptomic datasets (Fig. S9).

**Figure 6.**
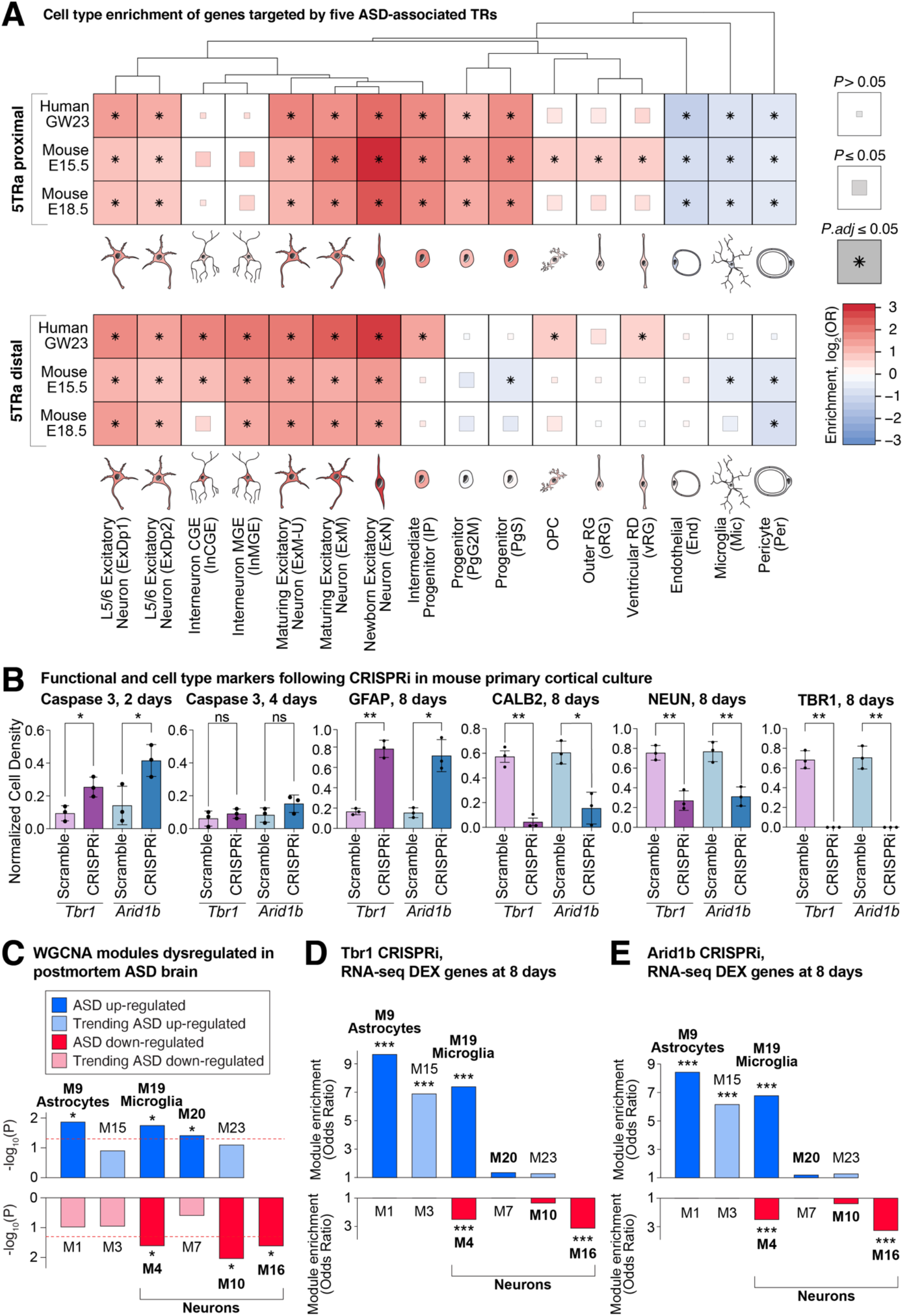
Cell type enrichment and functional consequences. **A)** Cell type clusters from the human fetal cortex (*33*) were assessed for enrichment of genes targeted by all five ASD-associated TRs and ATAC-seq (5TRa) in human (GW23) and mouse (E15.5 and E18.5) fetal cortex. The degree of enrichment is indicated by color; significance threshold is indicated by box size/asterisk. **B)** Cortical cells from postnatal day 0 dCAS9-KRAB mice were infected with lentiviral sgRNAs to the promoters of *Arid1b*, *Tbr1*, or scrambled controls. Immunohistochemistry was used to compare five markers. **C)** Down- (red) and up- (blue) regulated modules of co-expressed genes were previously identified in the prefrontal cortex of individuals with ASD (*7*). **D)** Differentially expressed genes, following CRISPRi to *Arid1b* were identified from bulk RNA-seq of day 8 cultured cells. The enrichment of differentially expressed genes is shown in the modules from ‘C’. **E)** The analysis in ‘D’ is repeated for CRISPRi to *Tbr1*. GW23: gestational week 23; E15.5/E18.5: embryonic day 15.5/18.5. Statistical analyses: A: Fisher’s exact test; B: Two-tailed T-Test with Tukey correction was used for pairwise comparisons; D-E: Fisher’s exact test; * ≤ 0.05, *** ≤0.001.

### Reduction of ARID1B and TBR1 expression in cultured neonatal mouse cortical cells reduces neuronal density and increases glial density, recapitulating human *postmortem* patterns of gene expression

The overlapping peaks of ASD-associated TRs suggests a convergent mechanism through which disruption of multiple genes can lead to similar neurodevelopmental phenotypes; this predicts similar functional consequences to changes of their expression. To test this prediction, we used lentivirus to deliver CRISPRi sgRNAs, designed to target proximal 5TRa loci in the promoter region of *Arid1b* or *Tbr1*, in primary cortical cultures collected from P0 dCAS9-KRAB mice. Compared to scrambled sgRNAs, CRISPRi reduced relative expression of *Arid1b* to 75% by day 2 and to 25% by day 8; likewise, *Tbr1* was reduced to 50% by day 2 and to 5% by day 8 (Fig. S10). CRISPRi to either *Arid1b* or *Tbr1* increased Caspase 3 on day 2, but not day 4, suggesting a transient increase in apoptosis. By day 8, immunohistochemistry showed there was a reduction of neuronal marker expression (∼3-fold reduction in NeuN^+^ cells and ∼6-fold reduction of CALB^+^ cells), accompanied by an increase in astrocyte marker expression (∼4.5-fold increase in GFAP^+^ cells; Fig. 6B).

Bulk RNA-seq analysis of the *Arid1b* and *Tbr1* CRISPRi-treated cells identified numerous differentially expressed genes (absolute fold-change ≥2, *P.adj* ≤ 0.05, Table S7). To assess how these results related to patterns of gene co-expression module dysregulation observed in the *postmortem* cortex of individuals with ASD (*7*), we considered the enrichment of these modules for the differentially expressed genes (Fig. 6C). Remarkably, both the *Arid1b* and *Tbr1* CRISPRi-treated cortical cells produced a pattern similar to that seen in ASD human brains. Neuronal modules that are down-regulated in the ASD brain (M4, M16) were enriched for down-regulated genes following CRISPRi (P ≤ 7×10^-9^, Fig. 6D-E, Table S7). On the other hand, astrocyte and glial modules that are up-regulated in the ASD brain (M9, M19) were enriched for up-regulated genes following CRISPRi (P ≤ 9×10^-12^, Fig. 6D-E, Table S7). While the direction of effect was similar, the magnitude differed between the cellular assay and *postmortem* brain data (*8*). Cell type deconvolution for six major cell types in *postmortem* human data suggests a 2.3% decrease in upper layer excitatory neurons in ASD compared to control (*P.adj* = 0.046, logistic regression with RIN, PMI, sex, and age as covariates, Fig. S9) with other cell types differing by 3.4% or less, in contrast to the dramatic changes in cell type composition in the CRISPRi primary cortical culture assay (Fig. 6B).

## Discussion

Over half the genes associated with ASD play a role in gene regulation, suggesting that transcriptional dysregulation is a major etiological factor in ASD (*2, 34, 35*). Here, we used ChIP-seq to identify regulatory targets of five of these ASD-associated TRs (ARID1B, BCL11A, FOXP1, TBR1, and TCF7L2) in the developing human and mouse cortex; we found that they converge on about 15,000 loci (6.5Mbp) proximal to the transcription start site of genes that are highly expressed in the developing brain, along with 5,000 distal loci (1.5Mbp). This overlap is surprising, since four of the five genes have a distinct DNA-binding domain and known motif (Fig. S4), they belong to different transcription regulator classes, and they have variable expression patterns across tissues and cell types. Despite this heterogeneity, we show that this overlap is greater than expected compared with ChIP-seq data from other TRs in heterogeneous tissues (Fig. 2A-B) and is also observed at two developmental stages in the mouse cortex (Fig. 2E-I).

Our results provide a parsimonious explanation for how disruption of multiple ASD-associated genes lead to a common diagnostic entity. Out of 102 ASD-associated genes (*2*), 101 have three or more of the ASD-associated TRs binding in open chromatin regions near their transcription start site (proximal ≥3TRa); 96 of these are targeted by all five ASD-associated TRs (proximal 5TRa, Fig. 4). This provides a mechanism by which disruption of each gene can impart risk to ASD and by which this risk can converge to a shared phenotype across many genes. We predict that other ASD-associated genes with a role in transcriptional regulation will also follow this pattern.

While the overlap of ASD-associated TR binding sites provides a potential mechanism for convergent ASD risk, it presents a challenge for explaining specificity to ASD, since almost half of all protein-coding genes share the proximal 5TRa pattern. Considering epigenetic and transcriptomic data from the developing human cortex provides a potential solution. The ASD-associated TR binding sites overlap substantially with loci identified by ATAC-seq (open chromatin) (*25*) and H3K27ac ChIP-seq but show minimal overlap with H3K27me3 ChIP-seq peaks (Fig. 2C-H); this suggests a predominant role in transcriptional activation. This conclusion is supported by the strong correlation between the number of ASD-associated TRs at proximal open chromatin sites and the level of gene expression in the developing human cortex (*6*). Analysis of single cell transcriptomic data from the developing human cortex (*17, 33, 36*) suggest that these observations arise from cells of the neuronal lineage (Fig. 6A). From these data, we might predict that the predominant role of these ASD-associated TRs is to increase gene expression during development of neuronal lineage cells. Since ASD genes are preferentially expressed in neuronal lineage cells (*2, 6, 19*), and are amongst the most highly expressed genes in the cortex (Fig. 4I), they may be especially vulnerable to perturbation of ASD-associated TRs. Binding of ASD- associated TRs at distal sites may further add to this vulnerability, as suggested by our analysis of VISTA element hs399 and its role in *BCL11A* expression (Fig. 5).

Protein-protein interactions between the TRs could explain the shared genomic targets (Fig. 2) and the reduced activity of the hs399 Vista enhancer in the absence of TBR1 (Fig. 5). Protein interaction data from the literature provide some support for this possibility, including interactions of BCL11A-TBR1 (*37*), FOXP1-TBR1 (*38*), FOXP1-BCL11A (via NR2F1/NR2F2) (*39*), FOXP1-TCF7L2 (*40*), and FOXP1-YY1 (Fig. 3) (*39*). Motif analysis results would also fit with this explanation. Four of the five ASD-associated TRs have previously been associated with a distinct DNA-binding motif based on DNA sequence enrichment within ChIP-seq peaks (Fig. S4). While we do observe enrichment of the BCL11A, FOXP1, TBR1, and TCF7L2 motifs (Fig. S4), the degree of enrichment is comparatively modest (Fig. 3, Table S4). In contrast, other DNA-binding motifs, such as the motif of the ASD-associated genes *RFX3* and *TCF4*, are substantially enriched (Fig. 3). These motif data also support the hypothesis that one of the core roles of these five TRs is to drive the expression of genes in neuronal lineage cells through binding to the promoter region. The observed motifs (Fig. 3) and hs399 Vista enhancer experiment (Fig. 5) implicate the formation of promoter-enhancer loops in this role, based on the enrichment for the GFY-STAF-ZNF143 proximal motif and CTCF/BORIS distal motif (Fig. 3). These patterns of protein-protein interactions may reflect the role of these ASD-associated TR proteins in known chromatin regulatory complexes, including BAF/SWI/SNF (ARID1B, BCL11A, TCF7L2) and NuRD (BCL11A, FOXP1) (*41–43*).

If regulatory ASD-associated genes act through complexes at promoter and/or enhancer regions to increase gene expression in neuronal lineage cells, we would predict decreased expression of these target genes would confer ASD risk. This fits conceptually with exome-sequencing results, which identified ASD risk through loss-of-function variants in genes with high neuronal expression (*2, 6*) and whole-genome sequencing results, which implicate *de novo* variants in promoter regions conserved across species (*44*). Regulatory complexes also provide a mechanism by which multiple common variants of small effect size could lead to additive risk (*45, 46*). Under this regulatory complex model, we would predict that heterozygous disruption of one of these ASD-associated TRs would lead to dysregulation of other ASD genes. This expectation is borne out in mice with heterozygous loss-of-function of *Foxp1* (*47*), however, the interpretation is challenging due to experimental heterogeneity, including differences in brain region, developmental stage, mutation, and the varying sensitivity to detect differential gene expression based on baseline levels of gene expression.

As a complementary approach to assessing the functional impact of loss-of-function of ASD-associated TRs, we used CRISPRi to knockdown *Arid1b* and *Tbr1* in primary cultures from P0 mouse neocortex. As predicted, we observed substantially decreased expression of numerous ASD-associated genes; corresponding increased expression was observed for genes in previously defined co-expression modules enriched for astrocytes and microglia marker genes (Fig. 6D-E). These results mirror patterns observed from bulk and single cell analyses of ASD cases and controls in the *postmortem* human brain (*7, 9, 48*). Concurrent analysis of cell type markers suggests that these signals are driven by changes in cell-type proportion, specifically a relative reduction in neurons and increase in astrocytes (Fig. 6B). A corresponding transient increase in Caspase 3 suggests increased apoptosis in the presence of the knockdown; thus, much of the observed transcriptional changes may reflect an alteration in the ratio of cell types.

Similar changes in cell-type proportion have been observed following disruption of cortical transcription factors, including *TBR1*, in human organoid models (*49*) and an increase in the proportion of protoplasmic astrocytes was described in the *postmortem* cortex of ASD cases (*9*). Deconvolution of bulk transcriptomic data from the *postmortem* brain in ASD cases and controls shows similar patterns, however the changes are modest and heterogenous (Fig. S9) (*8*).

Limitations to our analysis include the reliance on bulk cortical tissue to derive the ChIP-seq and ATAC-seq and peaks, so we cannot directly differentiate between cell types. However, some cell type information can be gleaned, since *BCL11A* and *TBR1* cortical expression appear to be specific to the neuronal lineage and *TBR1* is preferentially expressed in deep layer excitatory neurons (*20*), while *TCF7L2* is expressed in progenitor cells (*50*). None-the-less, we cannot be sure that the observed overlaps in TR binding sites exist in individual cells versus representing a pattern across multiple cell types. While the model of regulatory complexes bound to shared promoter and/or enhancer regions potentially explains convergence across TR ASD-associated genes, it is unclear how disruption of ASD-associated genes that are not thought to be TRs (e.g., *SCN2A*, *SYNGAP1*, *SLC6A1*) can lead to equivalent behavioral outcomes. However, convergence of ASD gene function in regulating synaptic function, as seen for TBR1 (*20, 51*) is a parsimonious explanation. Furthermore, while this regulatory complex model predicts widespread downregulation of highly brain-expressed genes, especially those associated with ASD, more data is required to verify this prediction. It is also unclear to what extent gene dysregulation or changes in the proportion of neuronal-lineage cell-types contribute to ASD symptomatology. Furthermore, it is unclear the extent to which a 75-95% knockdown using CRISPRi in cultured mouse cortical neurons should reflect a 50% constitutive knockout in developing human brain. These results raise a potential convergent downstream consequence of regulatory disruption, but substantial work is required to assess whether similar consequences occur in ASD or contribute to symptoms.

## Conclusions

Analysis of five ASD-associated transcription regulators leads to a model in which their encoded proteins act as components of molecular mechanisms titrated to control gene expression in developing neuronal lineage cells. Like a clock mechanism, many components are essential, and failure of any individual component can impact overall function. Under this model, disruption of any of multiple ASD-associated transcriptional regulator genes leads to a common neurodevelopmental outcome through shared genomic targets, while specificity to ASD and developmental delay is due to a combination of haploinsufficiency and high neuronal expression of ASD-associated target genes during neurodevelopment, making them the most vulnerable to small perturbations in expression.

## Acknowledgements

This work was supported by funding provided by the Simons Foundation Autism Research Initiative (SFARI), grants: 630332 (to J.L.R.R.) and 736613 (to S.J.S.), by the National Institute of Neurological Disorders and Stroke (NINDS), grant: R01 NS099099 (to J.L.R.R.), the National Institute of Mental Health (NIMH), grants: U01 MH116487 (to M.W.S.), U01 MH115747 (to M.W.S.), R01 MH129751 (to S.J.S.), R01 MH125516 (to S.J.S.), and U01 MH122681 (to S.J.S.), the Overlook International Fund, the National Research Foundation of Korea, grants: NRF-2020R1C1C1003426 (to J.Y.A.) and NRF-2021M3E5D9021878 (to J.Y.A.), and by the Korea University Insung Research Grant K2218731 (to J.Y.A.), and by a NSF graduate research fellowship (to N.F.P).

## STAR METHODS

### LEAD CONTACT AND MATERIALS AVAILABILITY

All unique/stable reagents generated in this study are available from the Lead Contact, Dr. John L. Rubenstein.

### EXPERIMENTAL MODEL AND SUBJECT DETAILS

#### Animals

All procedures and animal care were approved and performed in accordance with the University of California San Francisco Laboratory Animal Research Center (LARC) guidelines. All wildtype strains were maintained on a CD1 background. Animals were housed in a vivarium with a 12hr light, 12hr dark cycle. Postnatally, experimental animals were kept with their littermates. For timed pregnancies, noon on the day of the vaginal plug was counted as embryonic day 0.5.

### TRANSGENIC ANIMAL MODELS

dCAS9-KRAB mouse was a gift from McManus lab (UCSF) on a CD1 background. The dCas9-KRAB mice were generated in the FVB background with the TARGATTTM site-specific knock-in technology (*52*) by introducing a construct expressing containing CAG promoter, puromycin resistance, mCherry and the dead Cas9 (dCas9) protein fused to the KRAB (Krüppel Associated Box) domain into the *Hipp11* locus. hs399-CT2IG enhancer transgenic mouse was generated at the Gladstone Transgenic Gene Targeting Core.

### METHOD DETAILS

#### Transcription Regulator Chromatin Immunoprecipitation and Sequencing (TR ChIP-Seq)

Transcription regulator ChIP (TR ChIP) was performed using antibodies (specified below) against ARID1B, BCL11A, FOXP1, TBR1, and TCF7L2. Wildtype mouse cortices were dissected from E15.5 and E18.5 brains in ice-cold HBSS.

De-identified tissue samples were obtained with patient consent in strict observance of the legal and institutional ethical regulations. Protocols were approved by the Human Gamete, Embryo, and Stem Cell Research Committee, the institutional review board at the University of California San Francisco (UCSF). Fresh fetal brain samples were obtained from elective terminations, with no karyotype abnormalities or genetic conditions reported, and transported in freshly made Cerebral Spinal Fluid on ice (CSF). Samples were collected from gestational week 23 (GW23) prefrontal cortex (PFC). All dissections and ChIP-seq experiments were performed within two hours of tissue acquisition.

Human and mouse cortical samples were dissociated by pipetting in ice-cold HBSS. Dissociated cells were fixed in 1% formaldehyde for 10 min at RT. The fixed cells were neutralized with 1 mL 2.5M glycine and washed 3X in 1X PBS on ice. Fixed cells were lysed in a hypotonic buffer (50 mM Tris-HCl pH=7.5, 0.5% NP-40, 0.25% Sodium Deoxycholate, 0.1% SDS, 150 mM NaCl). Nuclei was extracted by centrifugation at 13,500 rpm for 10 min at 4°C and sheared into 200 - 1,000 bp fragments using a Covaris S2 (14 cycles of duty cycle = 5%, intensity = 3 and cycles per burst = 200).

Immunoprecipitation (IP) reactions of two biological replicates on mouse cortex at E15.5 and E18.5 and human PFC at GW23 were performed by diluting the sheared chromatin 1:10 in ChIP dilution buffer (16.7 mM Tris-HCl pH=8.0, 1.2 mM EDTA, 167 mM NaCl, 0.01% SDS, 1.1% Triton X-100) in 3 mL final volume. 100 mL was removed as “input”. 5 μg of primary antibody against ARID1B (Santa Cruz Biotech, sc32762 X), BCL11A (Abcam, ab19487), FOXP1 (Santa Cruz Biotech, sc-376650 X), TBR1 (Santa Cruz Biotech, sc48816 X) and TCF7L2 (Santa Cruz Biotech, sc166699 X) were added to each IP. 20X molar excess blocking peptide (FOXP1) and IgG (ARID1B, BCL11A, TCF7L2) and TBR1 constitutive null were used as negative control for each given ChIP experiments.

Antibody specificity has been examined for all the target proteins. The antibody specificity against TBR1 (*20*), BCL11A (*53*) and ARID1B (*54*) were previously demonstrated. We assessed anti-FOXP1 antibody specificity using a blocking peptide designed against the antibody epitope (Santa Cruz Biotech, sc-376650 P). TCF7L2 antibody specificity was examined through IHC analysis the TCF7L2 staining in the Tcf7l2^f/f^::1538CRE-ER::tdTomato^f/+^ conditional mutant compared to the heterozygous control (Tcf7l2^f/+^::1538CRE-ER::tdTomato^f/+^) at E12.5. Using this paradigm, we observed TCF7L2 signal reduced in pre-optic area (POA) in the conditional null mutant compared to the heterozygous controls, a region where CRE-expressing lineage cells are present.

Protein/antibody complexes were collected using Dynabeads (20 μL protein A + 20 μL protein G). Beads were washed once in each of Wash buffers (low salt buffer: 0.1% SDS, 1% Triton X-100, 2 mM EDTA, 20 mM Tris-HCl pH=8.0, 150 mM NaCl; High Salt buffer: 0.1% SDS, 1% Triton X-100, 2 mM EDTA, 20 mM Tris-HCl pH=8.0, 500 mM NaCl; LiCl buffer: 0.25 M LiCl, 1% IGEPAL CA630, 1% deoxycholic acid, 1 mM EDTA, 10 mM Tris-HCl pH=8.0 and lastly 1X TE pH=7.5. ChIP DNA was eluted twice in elution buffer (10 mM Tris-HCl pH=8.0, 1 mM EDTA, 100 mM NaHCO_3_, 1 mM SDS) at 65°C for 15 min each with shaking. Eluted ChIP DNA was reverse cross-linked in 8 μL 5M NaCl, 4 µL 1M Tris-HCl pH=6.5, 4 µL 0.5M EDTA overnight at 65°C. ChIP DNA was treated with 4 μL 10 mg/ml RNase A at 37°C for 15 min and then 1 µL 10 mg/mL proteinase K to each sample and incubated at 55°C for 1 hr.

ChIP DNA was purified using ChIP DNA Clean and Concentrator kit (Zymo Research, D5205). ChIP-seq libraries were generated using Ovation Ultralow System V2 Multiplex System (NuGEN) following manufacturer’s protocol, using 12 PCR cycles. The resulting libraries were size selected 180–350 bp using BluePippin (Sage Science) and sequenced at the Center for Advanced Technology at UCSF (Illumina HiSeq 4000; http://cat.ucsf.edu/) using a single read 50-bp strategy.

#### Histone Chromatin Immunoprecipitation and Sequencing (Histone ChIP-Seq)

Human and mouse samples were acquired as for TR ChIP-seq above. All dissections and downstream experiments were performed within two hours of tissue acquisition. From each dissection, nuclei were isolated by manually douncing the tissue twenty times in 1 mL Buffer I (300 mM sucrose, 60 mM KCl, 15 mM NaCl, 15 mM Tris-HCl pH=7.5, 5 mM MgCl_2_, 0.1 mM EGTA, 1 mM DTT, 1.1 mM PMSF, 50 mM Sodium Butyrate, EDTA-free Protease inhibitors) on ice using a loose pestle douncer, and then lysed on ice for 10 min after adding 1 mL Buffer II (300 mM sucrose, 60 mM KCl, 15 mM NaCl, 15 mM Tris-HCl pH=7.5, 5 mM MgCl_2_, 0.1 mM EGTA, 0.1% NP-40, 1 mM DTT, 1.1 mM PMSF, 50 mM Sodium Butyrate, EDTA-free Protease inhibitors).

During the incubation, nuclei were counted using trypan blue and 500,000 nuclei were spun down at 7,000rpm for ten minutes at 4°C. Nuclei were resuspended in 250 μL MNase buffer (320 mM sucrose, 50 mM Tris-HCl pH=7.5, 4 mM MgCl_2_, 1 mM CaCl_2_, 1.1 mM PMSF, 50 mM Sodium Butyrate) and incubated in a 37°C water bath with 2 μL MNase enzyme (NEB) for eight minutes. MNase digestion was stopped by adding 10 μL 0.5M EDTA, and chromatin was spun down for 10 min at 10,000 rpm 4°C. Soluble fraction S1 supernatant was saved at 4°C overnight, and S2 fraction was dialyzed overnight in 250uL dialysis buffer at 4C (1 mM Tris-HCl pH=7.5, 0.2 mM EDTA, 0.1 mM PMSF, 50 mM Sodium Butyrate, 1X Protease Inhibitors).

S1 and S2 fractions were combined, 50 μL was saved as input, and immunoprecipitation assay was set up in 50 mM Tris-HCl pH=7.5, 10 mM EDTA, 125 mM NaCl, 0.1% Tween-20. 250 mM Sodium Butyrate was supplemented for H3K27ac ChIPs. The following antibodies were used for ChIP: H3K27ac (Millipore, cma309), H3K4me1 (Abcam, ab8895), H3K27me3 (Millipore, 07-449), H3K4me3 (Abcam, ab185637). 1 microliter of antibody was added to 1 mL chromatin in ChIP dilution buffer (16.7 mM Tris-HCl pH=8.0, 1.2 mM EDTA, 167 mM NaCl, 0.01% SDS, 1.1% Triton X-100) and incubated overnight with chromatin at 4C rotating. Protein A and Protein G beads (10 mL each) were blocked overnight in 700 mL ChIP buffer, 20 mL yeast tRNA (20 mg/mL), and 300 mL BSA (10 mg/mL). Beads were washed three times on ice in Wash buffer I (50 mM Tris-HCl pH=7.5, 10 mM EDTA, 125 mM NaCl, 0.1% Tween-20, supplemented with 1X protease inhibitors and 5 mM sodium butyrate) and three times in Wash buffer II (50 mM Tris-HCl pH=7.5, 10 mM EDTA, 175 mM NaCl, 0.1% NP-40, supplemented with 1X protease inhibitors and 5 mM sodium butyrate). Lastly, beads were washed once in 1X TE buffer. ChIP DNA was eluted twice in 100 mL elution buffer (10 mM Tris-HCl pH=8.0, 1 mM EDTA, 100 mM NaHCO_3_, 1 mM SDS) at 37°C for 15 min each with shaking.

IP reactions were incubated at 65°C for 30 min and purified using ChIP DNA Clean and Concentrator kit (Zymo Research, D5205). ChIP-seq libraries were generated using Ovation Ultralow System V2 (NuGEN) following manufacturer’s protocol. The resulting libraries were size selected (180–350 bp) and sequenced at the Center for Advanced Technology at UCSF (Illumina HiSeq 4000; http://cat.ucsf.edu/) using a single read 50-bp strategy.

#### ChIP-seq Computational Analysis

##### Peak Calling

Human samples were aligned to the main chromosomal contigs of the GRCh38 genome. Mouse samples were aligned to the main chromosomal contigs GRCm38 (mm10) genome. Both alignments were performed by BWA (v0.7.15) *bwa mem ref_genome.fa sample.fastq.gz*. Resulting SAM files were converted to BAM files with a MAPQ filter of 30 and sorted using samtools (v1.10). *samtools view -q 30 -Shu -o sample.unsorted.bam sample.unsorted.sam*, and *samtools sort -o sample.bam sample.unsorted.bam*.

For all phenotypes, significant peaks were identified against matched input controls or WT background using mac2 (v2.2.7.1). Narrow peak calling was used with a q-value cut-off of 0.01. Model-based peak calling and local significance testing were disabled. A fixed fragment extension length of 200bps was used. *macs2 callpeak -t sample.bam -c input.sample.bam -f BAM -g mm/hs -nolambda -nomodel -ext 200 -bdg -q 0.01*.

For mouse samples, biological replicates were kept separate when first identifying peaks. IDR analysis was used to confirm the quality of peaks between biological replicates. Aligned reads of the replicates were then merged and new peaks were called. The peaks derived from merged biological replicates were used for downstream analysis.

##### Coverage Heatmaps

Coverage heatmaps for transcription factor ChIPseq samples were generated using deepTools (v3.5.1). Biological replicates were kept separate in these heatmaps to better display coverage. The regions shown were pooled from peaks from every transcription factor of the designated age. The viewing reference point was set to *center* and a viewing range of 1kb was used. *computeMatrix reference-point -referencePoint center -S [sample1.bigwig, sample2.bigwig, … sampleN.bigwig] -R E15_peaks.bed -o E15_matrix.txt -a 1000 -b 1000.* Followed by, *plotHeatmap -m E15_matirx.txt -o E15_heatmap.pdf -missingDataColor white*.

##### Peak Annotation

Called peaks from the Chip-seq datasets were annotated to all transcripts on GENCODE version 31 (human; hg38) and GENCODE version 23 (mouse; mm10). Proximal peaks were defined as being 2,000bp upstream of the TSS. Where multiple TSS were present for a given gene, we used the union of all proximal regions. BedTools intersect was used to identify overlap between peaks and gene promoters (*55*) with any overlap between the ChIP-seq peak and promoter region defining the peak as “proximal”. Chip-seq peaks that did not overlap with promoter regions were defined as “distal” and the nearest TSS, identified by BedTools closest, was used to define the gene associated with distal peaks.

#### ENCODE ChIP-seq overlap analysis

Where replicate samples were present the replicate with the highest number of total peaks was selected (ENCODE references: ENCFF002EXB, ENCFF132PDR, ENCFF765EAP, ENCFF039EYW, ENCFF366KUG, ENCFF215GBK, ENCFF934JOM, ENCFF264HRE, ENCFF643ZXX, ENCFF390HDY, ENCFF551VXN, ENCFF996EBR, ENCFF634YGY, ENCFF751TBY). FASTQ files were processed to generate peaks using the methods described above (‘*Peak Calling’*). Similar numbers of total proximal and distal peaks were present in the ENCODE liver and ASD-associated TR cortex samples (21,295 proximal peaks in liver vs. 23,668 in cortex, P=0.61 Wilcoxon; 52,899 distal peaks in liver vs. 45,429 in cortex, P=0.61 Wilcoxon). Peaks lists were sorted by p-value, followed by genomic location (if duplicate p-values were present) and the top 10,000 proximal peaks (selected due to 10,746 proximal peaks in ZBTB33) and 8,000 distal peaks (selected due to 8,002 distal peaks in FOXP1) were selected to exclude peak count as a variable. Bedtools intersect (*55*) was used to identify the number of peaks at the same genomic location (any overlap was counted due to the large search space and to minimize the impact of peak size), expressed as a percentage of the peaks used (10,000 or 8,000). For peaks that did intersect, the correlation between the p-value and genomic location ranked lists was used to estimate Spearman’s rho (*23*). Similar patterns of overlap were observed when reducing the number of peaks used to 4,000 for both proximal and distal loci.

#### Motifs

Enriched known motifs were found using Homer (v4.10.3). A set window size of 200 and the repeated-masked sequence options were used. *findMotifsGenome.pl sample_peaks.narrowPeak hg38/mm10 motif.out -size 200 -mask -gc.* The output table *knownResults.txt* was used to generate the motif enrichment plot (Fig. 3).

#### Vista enhancer validation of distal ≥3TRa peaks

To determine whether ≥3TRa peaks are functional enhancers within the developing brain, we calculated the odds ratio of a locus overlapping a validated VISTA enhancer based on the presence or absence of a distal ≥3TRa peak. A bedtools intersect was performed on the list of 1,942 human VISTA enhancer regions and 13,875 distal ≥3TRa peaks. The VISTA enhancer atlas records the activity of each region across over 20 different tissues. The odds of being a positive VISTA enhancer in a given tissue based on the presence of a distal ≥3TRa peak was assessed via a two-sided Fisher’s exact test. Tissues with fewer than 50 positive VISTA enhancers were excluded from the analysis due to low sample size. The presence of a distal ≥3TRa peak significantly increased the odds of being a validated forebrain (OR=2.17, FDR=5.5 ×10^-5^, Table S6) or neural tube (OR=2.27, FDR=5.0 ×10^-4^, Table S6) enhancer (Figure S8A). A subset of VISTA forebrain enhancer sequences were retested *in vivo* and annotated by an expert developmental neuroanatomist for specific expression in the pallium, subpallium, or both and the analysis was repeated focusing on this subset alone (Figure S8B).

#### Identification of distal TR interactions with ASD genes via ABC score

The Activity-by-Contact (ABC) model identifies enhancer-gene relationships based on chromatin state and conformation and is more effective than distance-based approaches (*29*). Gestational week 18 (GW18) bulk ATAC-seq and H3K27ac ChIP-seq data from human fetal prefrontal cortex (*25*) were aligned to hg19 using the standard Encode Consortium ATAC-seq and ChIP-seq pipelines respectively with default settings and pseudo replicate generation turned off (https://github.com/ENCODE-DCC). HiC contacts with 10kb resolution from human GW17-18 fronto-parietal cortex were obtained from http://resource.psychencode.org/Datasets/Pipeline/HiC_matrices/PIP-01_DLPFC.10kb.txt.tar.gz (*30*) in an hdf5 format separated by chromosome. Hdf5 files were filtered for contacts with a score > 0 and converted into a bedpe format. Trimmed, sorted, duplicate and chrM removed ATAC-seq, sorted, duplicate removed ChIP-seq bam files, and HiC bedpe files from GW17-18 cortex were provided as input for calculating ABC scores.

According to the ABC score pipeline, (https://github.com/broadinstitute/ABC-Enhancer-Gene-Prediction) ATAC-seq and H3K27ac ChIP-seq bam files were provided as input to the MakeCandidateRegions.py script with the flags --peakExtendFromSummit 250 --nStrongestPeaks 150000. Candidate enhancer regions identified were then provided to the run.neighborhoods.py script in addition to hg19 transcript bounds merged by overlapping 2000bp promoters (Gencode v38lift37, basic). Finally, predict.py was used to identify final candidate enhancers using HiC data with the flags --hic_type bedpe --hic_resolution 10000 --scale_hic_using_powerlaw --threshold .02 --make_all_putative. All other settings for the ABC score pipeline remained constant.

To compare nearest neighbor vs ABC approaches for identifying enhancer-gene pairs, we identified the nearest protein coding gene (Gencode v31, all) to each of the 13,875 identified distal ≥3TRa peaks (ARID1B, BCL11A, TBR1, and ATAC-seq) using bedtools closest with the options “-d - t first”. Candidate enhancer-gene pairs from ABC score were converted to hg38 using UCSC liftOver and liftOverBedpe (https://github.com/dphansti/liftOverBedpe). Overlaps between ≥3TRa peaks and candidate enhancer-gene pairs were identified using bedtools intersect. In total, 269 distal ≥3TRa peaks were nearest neighbors to 54 unique ASD genes, and 285 peaks had ABC contacts to 77 ASD genes.

#### *In vitro* CRISPRi Assay in Mouse Primary Cortical Cultures

##### Guide Design

To generate lentiviral guide constructs against Transcriptional Start Site (TSS) of *Arid1b* and *Tbr1* using 2 kb upstream of TSS for each given gene. In addition to TSS-sgRNA, lentiviral guide constructs were also designed against putative regulatory elements (pREs), each of which are bound *in vivo* by all hcASD-TRs, forming “hubs”. The DNA sequences were inputted into CRISPOR tool (*58*) from Zhang lab at Massachusetts Institute of Technology (https://zlab.bio/guide-design-resources). To facilitate the cloning process, TTGG and CAAA were added to 5’ and 3’-ends of forward and reverse guides, respectively. Scrambled guides were designed using GenScript online browser tool from *Arid1b* and *Tbr1* TSS-sgRNAs (https://www.genscript.com/tools/create-scrambled-sequence). The complete list of guides and the corresponding genomic regions are shown below.

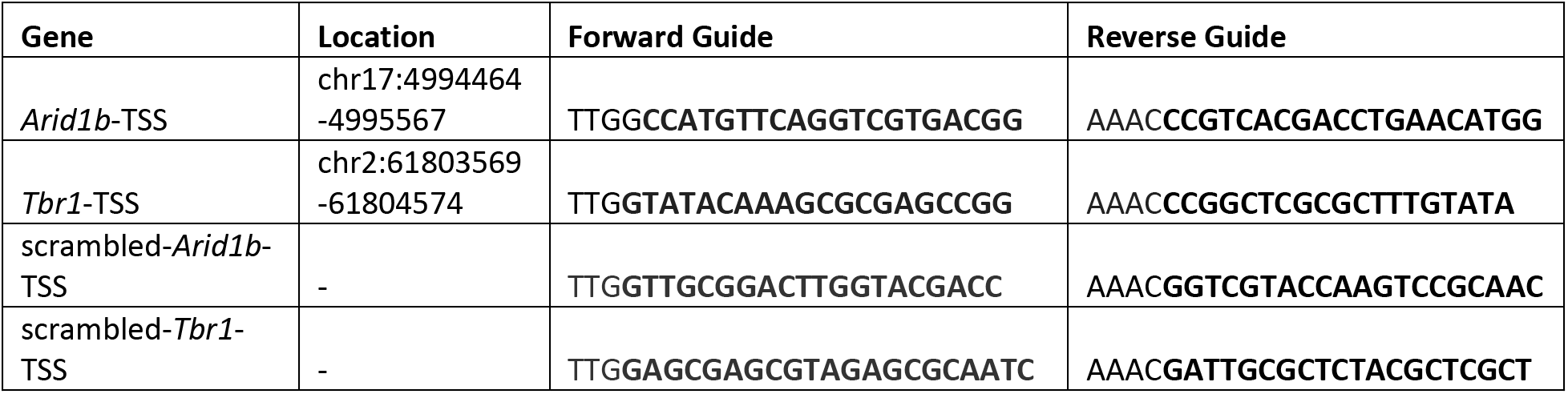

##### Generating sgRNA Lentivirus

Guide RNA oligonucleotides were annealed in 1X annealing buffer (100 mM Tris pH7.5, 1 M NaCl, 10 mM EDTA) by heating for 5 minutes at 95°C, then cooling down gradually to 25°C, 5°C/min. *U6-stuffer-longTracer-GFP* lentivirus vector was digested with AarI enzyme overnight and gel purified. The annealed guides were cloned into the digested *U6-stuffer-longTracer-GFP* lentivirus vector overnight at 16°C. Ligated guides were cloned into *Stbl3* cells (Thermofisher) and verified by sequencing at ElimBio using Elim Primer# 258124.

Upon sequencing validation, the gRNA lentivirus was generated in HEK293T cells by transfecting 3’10 cm dishes with 3 ug sgRNA plasmid, 1.5 ug psPAX2 packaging vector (Addgene), 1.5 ug pmD2G envelope vector (Addgene), 850 uL jetPRIME buffer and 18 uL jetPRIME. Three days post-transfection, the sgRNA expressing lentivirus was purified as described previously (*59*). Empty *U6-stuffer-longTracer-GFP* lentivirus vector was used to generate mock control lentivirus.

##### Primary Cell Culture and In Vitro CRISPRi Assay

Cortex was dissected from P0 dCAS9-KRAB pups and dissociated using papain dissociation kit following manufacturer’s protocol (Worthington). A total of 400,000 cells were seeded into 24-well tissue culture dishes containing fresh N5 medium (500 uL N2 supplement, 121 uL BPE (bovine pituitary extract), 10 uL 100 ng/uL FGF, 10 uL 100 ng/uL EGF, 5 mL FBS, 0.5 mL Pen/Strep in 50 mL DMEM) that were pre-coated with poly-L-lysine (10 mg/ml, Sigma) and then laminin (5 mg/ml, Sigma). Polybrene (Thermofisher) was added to each tube to facilitate transduction at a final concentration of 8 μg/ml. Concomitantly, the P0 dCAS9-KRAB cells were transduced by adding 20 uL concentrated virus at the time of seeding. The cultures were grown for 16 days *in vitro* and N5 media was replaced every 48 hrs. This experiment was repeated twice (n=2).

#### RNA extraction and cDNA synthesis

Total RNA was extracted from the primary cortical cultures at 2, 4, 8, 12 and 16 days-post-transduction (DPT) using RNeasy Plus^®^ Micro Kit (QIAGEN, Cat# 74034) following the manufacturer’s protocol. First strand cDNA was synthesized from 0.5 ug of total RNA using Superscript reverse transcriptase III following manufacturer’s protocol (Thermofisher).

#### Quantitative real time PCR (qPCR)

Quantitative real time PCR (qPCR) was performed to measure RNA levels using SYBR Green (Bio-Rad) and 7900HT Fast Real-Time PCR System. Gene-specific primers for exon #1 of *Bcl11a, Drd1, Hbb, ef1α* as well as *Gapdh* housekeeping genes (HKG) were designed using the Primer 3 program. The expression levels of the genes in mock control and each TSS-CRISPRi RNA were normalized to the expression levels of the HKGs. Subsequently, the gene expression levels in TSS-CRISPRi RNAs were measured relative to the mock control using DDCT method as previously described (*60, 61*) and averaged across 3 experimental replicates for each biological set (n=2 biological replicates).

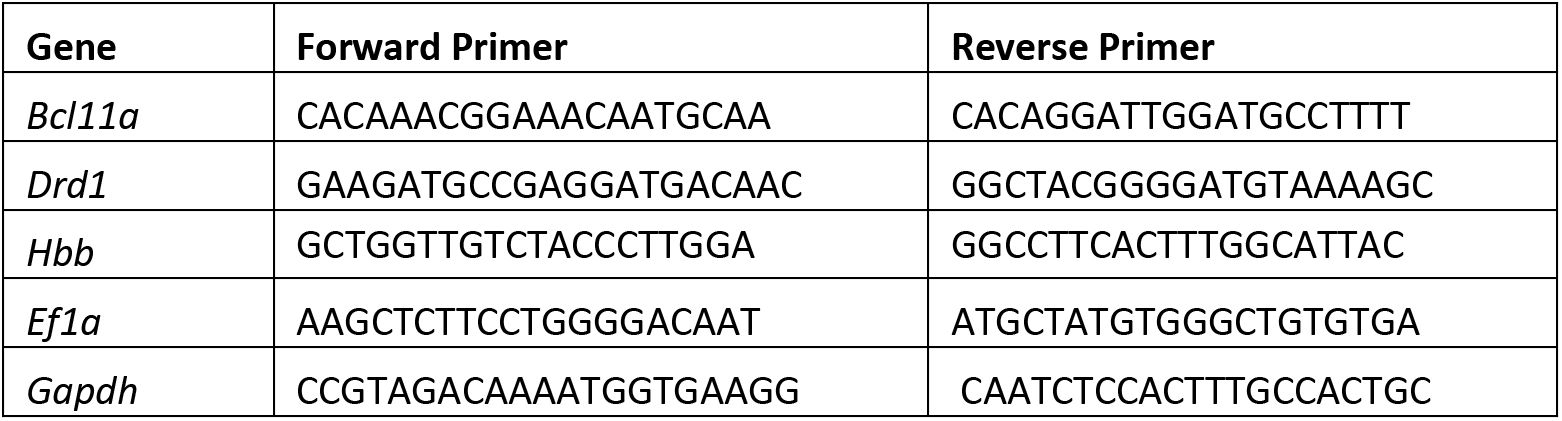

#### Generation of hs399-CT2IG Enhancer Transgenic Mouse

Enhancer hs399 was amplified from human genomic DNA and subcloned into Hsp68-CreERT2-IRES-GFP (*62*). Stable transgenic mice were generated by pronuclear injection at the Gladstone Transgenic Gene Targeting Core using the FVB strain. Founders were screened by PCR (*32*).

#### RNA-seq on TSS-CRISPRi Cells

Transcriptome profiling was conducted by using RNA-seq on *in vitro* dCAS9-KRAB (CRISPRi) cells 8 days-post-transduction by lentivirus encoding sgRNA guides against TSS of *Arid1b* and *Tbr1*. Approximately 300,000 cells were collected from each sample and immediately proceeded with RNA extraction using RNeasy^®^ Plus Micro Kit (QIAGEN) following manufacturer’s protocol. RNA quality was assessed using Agilent RNA 6000 Nano Kit (Agilent Technologies) and ran on Bioanalyzer 2100 (Agilent Technologies). Samples that had RIN scores of 8.5-10 were used to generate libraries. Library preparation and amplification was performed by TruSeq^®^ Stranded Total RNA Library Prep Kit with Ribo-Zero Gold Set A (Illumina, Cat# RS-122-2001). The amplification of adapter-ligated fragments was carried out for 12 cycles during which individual index sequences were added to each distinct sample. Library concentration was assessed with Qubit (Thermofisher, Cat# Q33231) and library fragment size distribution was assessed on the Agilent Bioanalyzer 2100 (Agilent Technologies) and Agilent High Sensitivity DNA Kit (Agilent Technologies) following manufacturer’s protocol. Pooled, indexed RNA-seq libraries were sequenced on HiSeq 4000 at Center for Advanced Technology (Illumina HiSeq 4000; http://cat.ucsf.edu/) to produce 150 bp paired-end reads.

Read count and transcript per million reads mapped (TPM) were determined using Salmon software version 1.3.0. A reference genome index for Salmon was created according to developer’s instructions for the mouse reference transcriptome from the GENCODE v23. Reads mapping and quantitation was simultaneously performed to individual transcripts. Gene-level counts were collated and normalized using the tximport and DESeq2 R Bioconductor packages (*63, 64*). We excluded the genes with zero count and selected the genes used in the enrichment analysis. Differential gene expression (DGE) analyses were performed on two replicates of TR TSS-CRISPRi against five control libraries, with following: ∼ replicate + condition. With random selections of control libraries, we did confirm our DGE consistently generate fold changes and statistics regardless of control sample batches. For gene set enrichment analysis, we used differentially expressed (DEX) genes of TSS-CRISPRi as those having adjusted p-value ≤ 0.05 and fold-change greater or less than 2, compared to control libraries. We collected a range of gene lists for ASD neurobiology and cortical development (*7, 33, 36*). DEX genes were converted to human ortholog genes using the HGNC annotation. Gene enrichment tests were assessed with a gene list using Fisher’s exact test with Bonferroni correction.

## Supplemental tables

TableS1_Peaks_Cortex_Submitted.xlsx: MACS2 output for the ChIP-seq and ATAC-seq data for human and mouse cortex

TableS2_Peaks_Liver_Submitted.xlsx: MACS2 output for the ChIP-seq data for human liver from ENCODE

TableS3_5TRa_Peaks_Submitted.xlsx: A list of all 5TRa loci in human GW23, mouse E15.5, mouse E18.5

TableS4_Motifs_Submitted.xlsx: Results of HOMER enrichment analysis for human GW23, mouse E15.5, mouse E18.5

TableS5_ASD_Genes_Submitted.xlsx: Types of peaks in proximity to 102 ASD-associated genes

TableS6_VISTA_TRs_Submitted.xlsx: Overlap between ChIP-seq data and VISTA data

TableS7_SingleCell_CRISPRi_Submitted.xlsx: 5TRa single cell enrichment and CRISPRi RNA-seq results

## Supplemental figures

**Figure S1.**
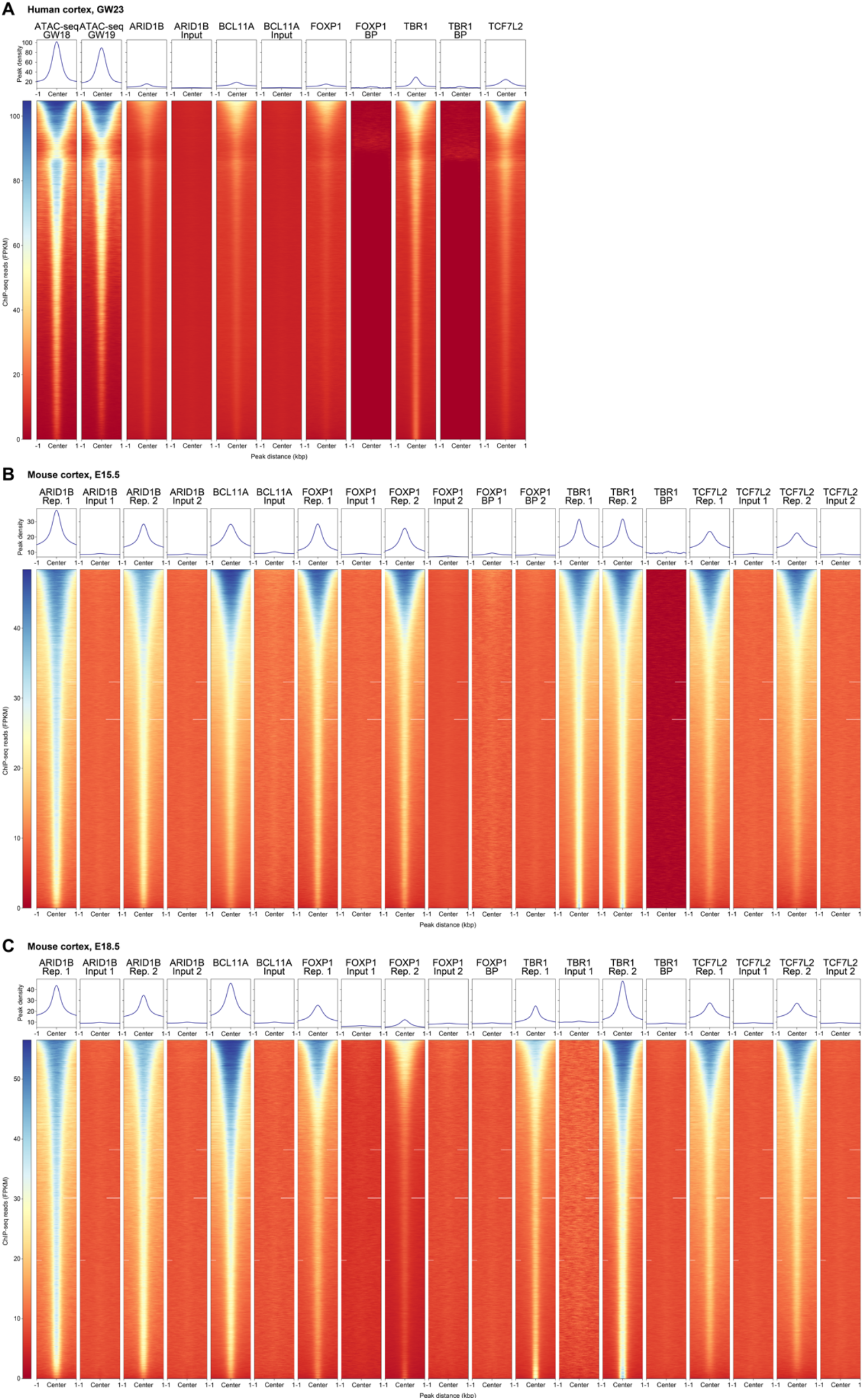
Read counts across all peaks in human and mouse. **A)** The analysis in Fig. 1D showing read counts with 1kbp around the middle of ChIP-seq peaks in human cortex but also including all peaks from ATAC-seq in human cortex performed at GW18 and GW19. The peaks are shown in the same order on the y-axis for all eight datasets, including inputs and blocking peptides (BP) as negative controls. **B)** Read counts arranged by peak count are shown for all replicates, inputs, and blocking peptides of ChIP-seq performed on mouse cortex at E15.5. Again, the peaks are shown in the same order on the y-axis for all datasets shown. **C)** The analysis in ‘B’ is repeated for all samples at E18.5 in mouse cortex.

**Figure S2.**
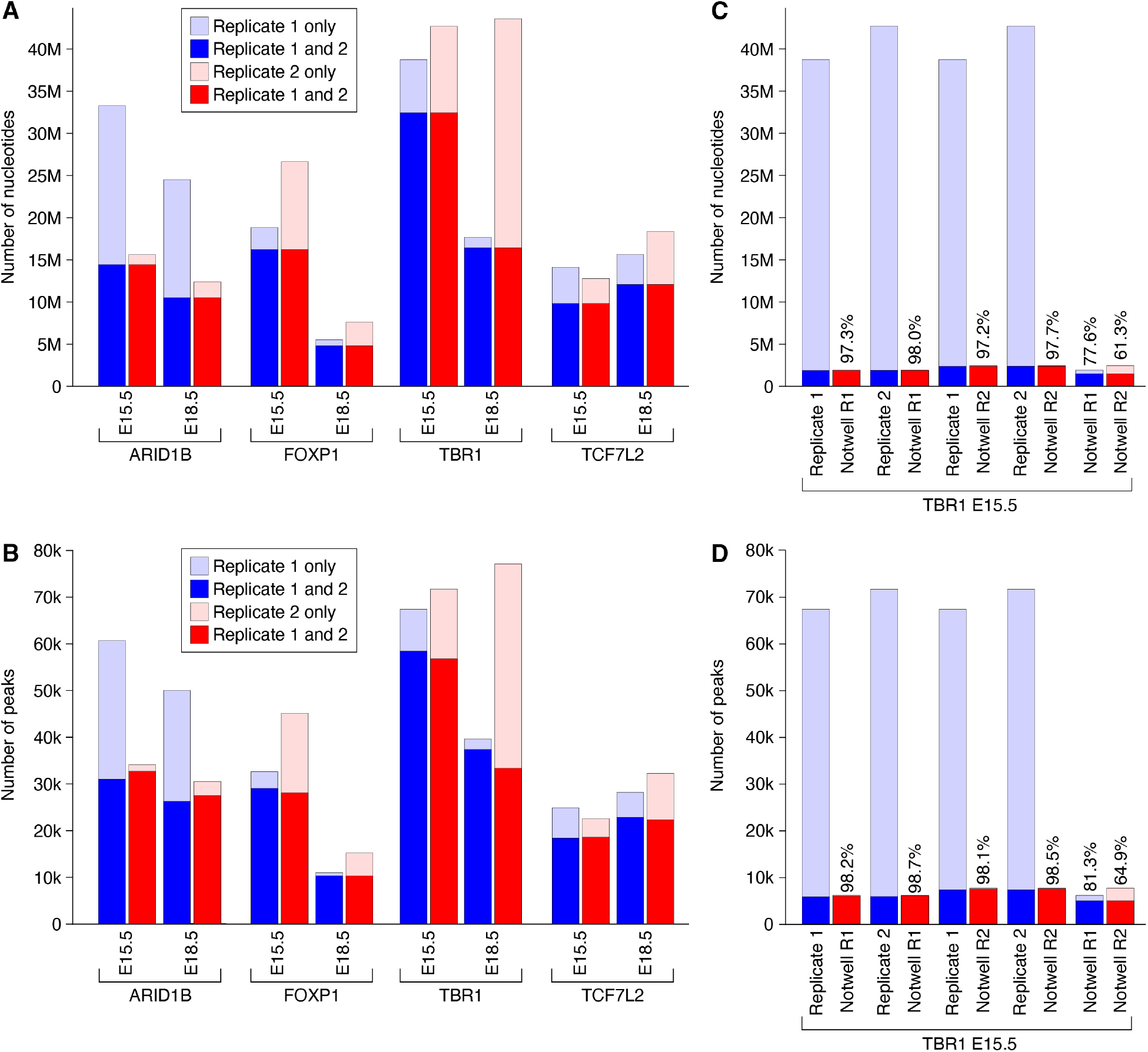
Reproducibility of peaks across replicates in mouse cortex. **A)** Two ChIP-seq replicates (replicate 1 in blue and replicate 2 in red) were performed for the four TRs (ARID1B, FOXP1, TBR1, and TCF7L2) at two developmental stages (E15.5 and E18.5). The number of nucleotides in each replicate is shown on the y-axis with darker colors showing nucleotides overlapping between replicates and lighter colors showing nucleotides unique to one replicate. **B)** The analysis in ‘A’ is repeated but with nucleotide count replaced by the number of peaks on the y-axis. The number of overlapping peaks (defined as any overlap) varies between replicates because one peak in replicate 1 may overlap multiple peaks in replicate 2 and *vice versa*. **C)** Using the same methods as ‘A’, TBR1 E15.5 Replicates 1 and 2 from this manuscript are compared to two biological replicates (Notwell R1 and Notwell R2) of TBR1 E15.5 mouse cortex data published previously (*13*). The percent of overlapping nucleotides is shown for the Notwell replicates. **D)** The analysis in ‘C’ is repeated but using overlapping peaks instead of overlapping nucleotides.

**Figure S3.**
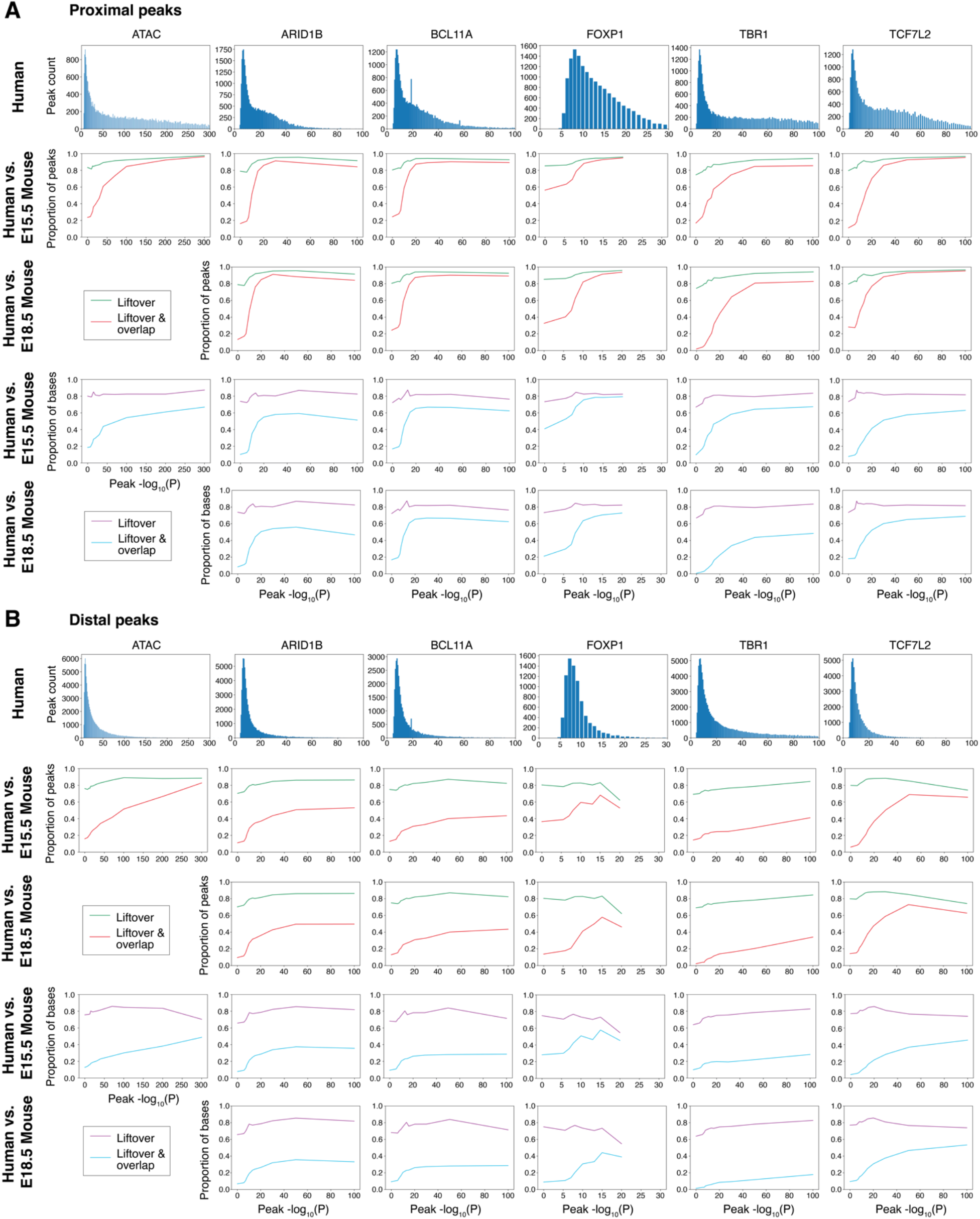
Peak conservation across species. **A)** First row: A histogram shows the distribution of ATAC-seq (GW18/19) and ChIP-seq (GW23) peaks from human cortex by MACS -log_10_(P) values at proximal loci. Second row: The human hg38 peaks were lifted over to mouse mm10 and compared to corresponding ATAC-seq and ChIP-seq data generated from E15.5 mouse cortex. The green line shows the proportion of human peaks that were successfully lifted over (1% minimum overlap) and the red line shows the proportion of human peaks that were successfully lifted over and had any degree of overlap with at least one mouse peak. Third row: The ‘second row’ analysis of repeated for ChIP-seq data generated from E18.5 mouse cortex. Fourth row: The ‘second row’ analysis is repeated but showing the proportion of human bases that were lifted over (purple) and lifted over and overlapping with a mouse peak from E15.5 data (cyan). Fifth row: The ‘fourth row’ analysis is repeated for data generated from E18.5 mouse cortex. **B)** The analysis in ‘A’ is repeated for distal peaks.

**Figure S4.**
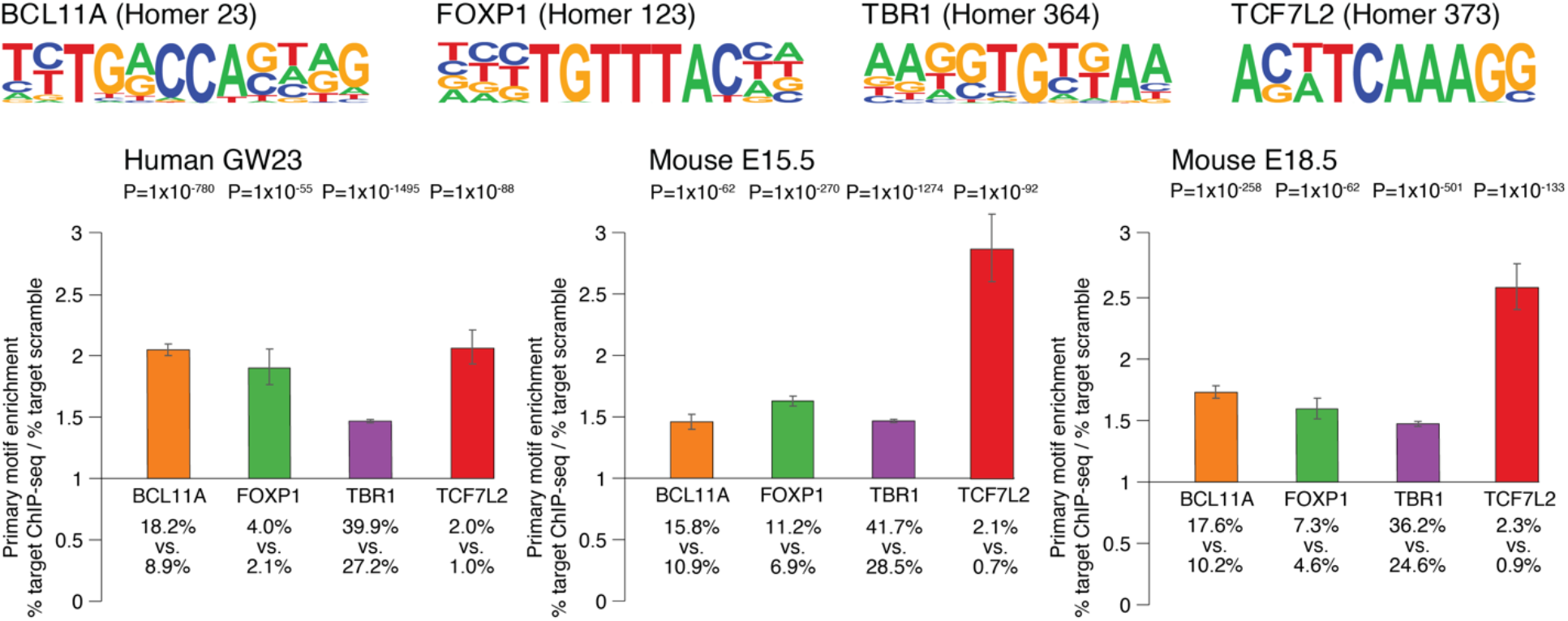
Enrichment of TR motif in ChIP-seq data. Motifs were identified from Homer for BCL11A, FOXP1, TBR1, and TCF7L2; no equivalent motif was available for ARID1B. Using Homer, the proportion of peaks with the corresponding motif (e.g., BCL11A motif in BCL11A ChIP-seq peaks) was compared to the proportion of sequence scrambled peaks with the corresponding motif to generate a relative risk (y-axis). Error bars represent the 95% confidence interval calculated from the log-transformed risk ratio; P-values are calculated by Homer using the binomial exact test.

**Figure S5.**
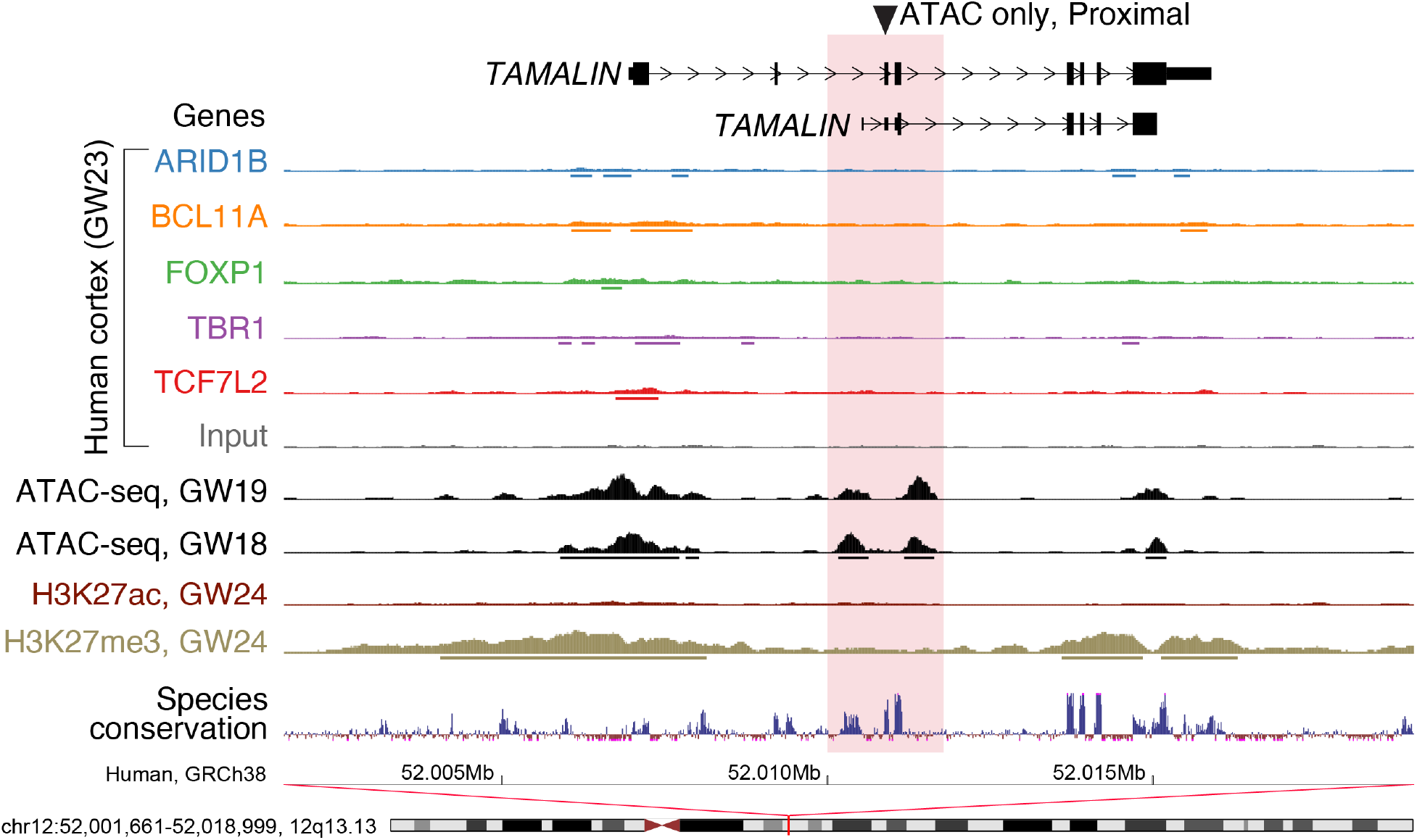
Region with ATAC-seq peak without TR ChIP-seq peaks. A representative example of ATAC-seq in GW18 and GW19 human prefrontal cortex and histone ChIP-seq data (H3K27ac, H3K27me3) in GW24 human prefrontal cortex without corresponding peaks from TR ChIP-seq data. Abbreviations: GW: gestation week.

**Figure S6.**
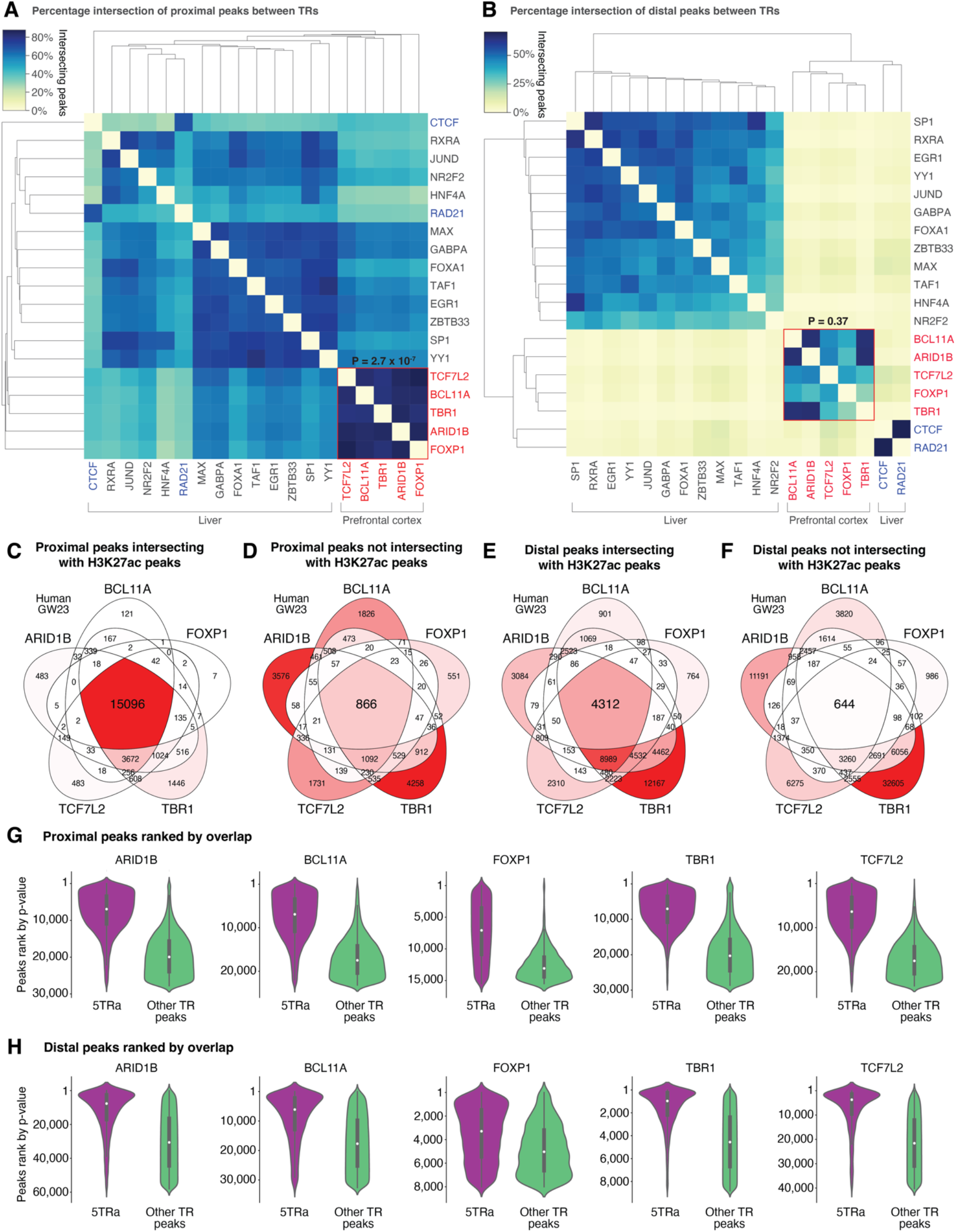
Overlap between TRs and epigenetic markers. **A)** Top 10,000 proximal ChIP-seq peaks ranked by p-value from 14 TRs in adult human liver (ENCODE) and five ASD-associated TRs in fetal human cortex at GW23 to assess the proportion of peaks that intersect. **B)** Equivalent plot for 8,000 distal peaks. **C)** Overlap between proximal ChIP-seq peaks for the five ASD-associated TRs in developing human cortex overlapping with H3K27ac ChIP-seq peaks. Color gradient represents the percentage of peaks in each section with red being the highest percentage and white being 0%; peak counts shown in each section. **D)** Equivalent plot for proximal peaks that do not overlap with H3K27ac ChIP-seq peaks, **E)** distal peaks that do overlap with H3K27ac ChIP-seq peaks, and **F)** distal peaks that do not overlap with H3K27ac ChIP-seq peaks. **G)** For each ASD-associated TR, ChIP-seq proximal peaks are ranked by p-value (with 1 being the highest confidence) and split between 5TRa peaks (left, purple) and all other TR peaks (right, green). H) Equivalent plot to ‘G’ for distal peaks. Abbreviations: 5TRa: peak with all five ASD-associated transcription factors (5TR) and ATAC-seq (a); GW23: gestational week 23. Statistical analyses: A and B: Wilcoxon test.

**Figure S7.**
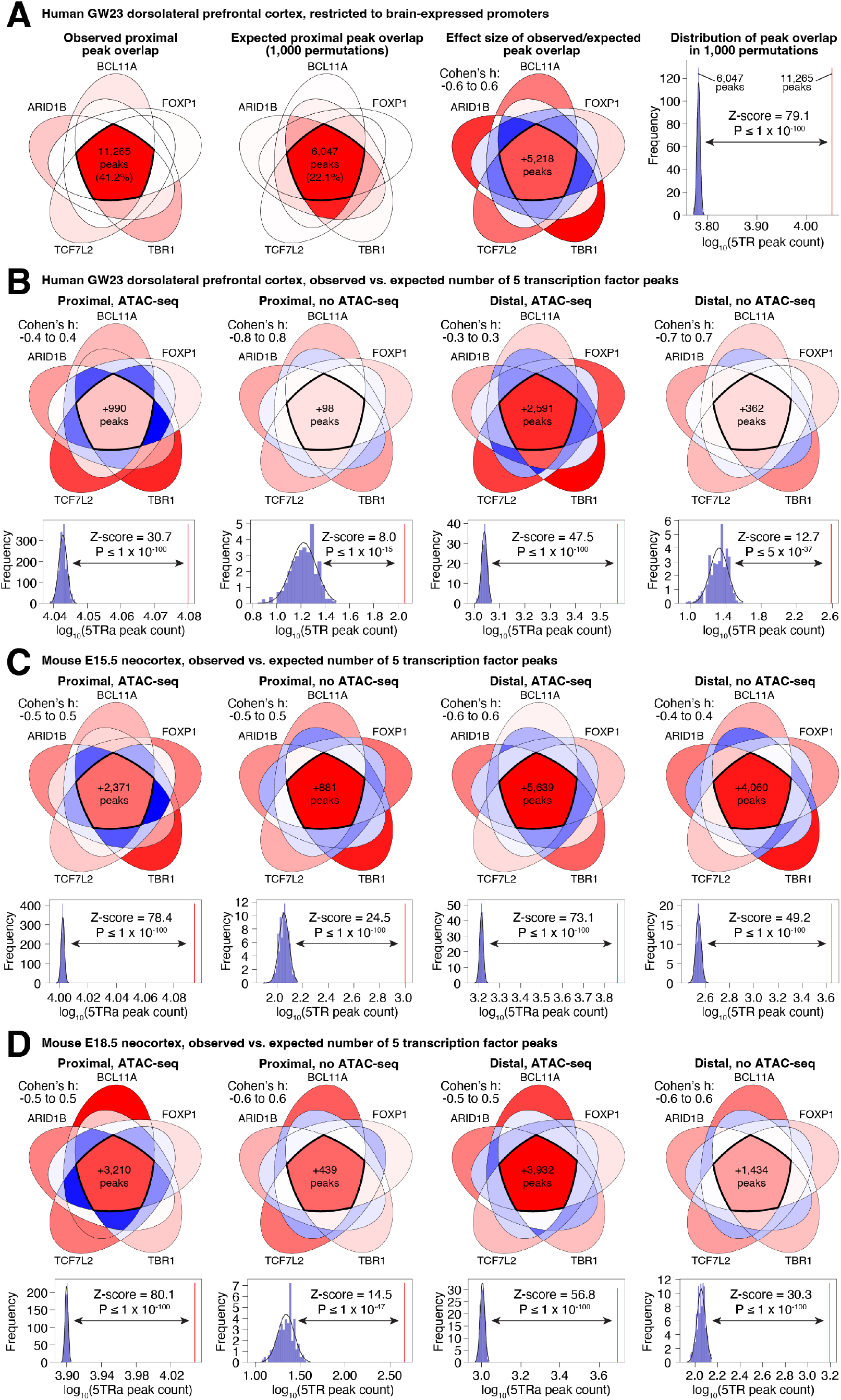
Permutation analysis of overlapping peaks. A) Left: Overlap of proximal peaks from each of the five ASD-associated TRs in GW23 human cortex was assessed and plotted as a Venn diagram. Intensity of red indicates the proportion of the total unique loci that are within that section, scaled from 0% (white) to the maximum (41.2%, red). **Middle left:** The expected number of overlaps is estimated by permutation analysis. For each permutation, the peaks of all five TRs are concatenated to create the search space, then, for each of the five TRs in turn, the same number of peaks as the original data are selected at random from the concatenated peak list. This is repeated for each of the five TRs then the overlap between permuted TRs is assessed. This process was repeated 1,000 times. The mean counts are shown in each section of the Venn diagram, with intensity of red scaled from 0% (white) to the maximum (22.1%, red). **Middle right:** The difference between the observed and mean permuted expectation and the standard deviation of the permutations is used to estimate the effect size (Cohen’s *d*) in each section. A positive Cohen’s *d* (more peaks in observed than expected) is shown in red, a Cohen’s *d* of 0 is white, while a negative Cohen’s *d* (fewer peaks in observed than expected) is shown in blue. Intensity of color represents a linear scale of Cohen’s *d* between -0.6 and 0.6. **Right:** The distribution of expected counts for the overlap of all five TRs (5TR) is shown as a histogram on a logarithmic axis with the observed count shown as a red line. The standard deviation is used to estimate the z-score, which is converted to a P-value. **B)** The methodology described in ’A’ is repeated for proximal (left) and distal (right) peaks, split by whether the peaks overlap with peaks from GW18/19 human cortex ATAC-seq. The effect size Venn diagram and histogram of expected vs. observed overlap of all five TRs are shown. **C)** The analysis in ‘B’ is repeated for E15.5 mouse cortex ChIP-seq and E15.5/E18.5 mouse cortex ATAC-seq data. **D)** The analysis in ‘B’ is repeated for E18.5 mouse cortex ChIP-seq and E15.5/E18.5 mouse cortex ATAC-seq data.

**Figure S8.**
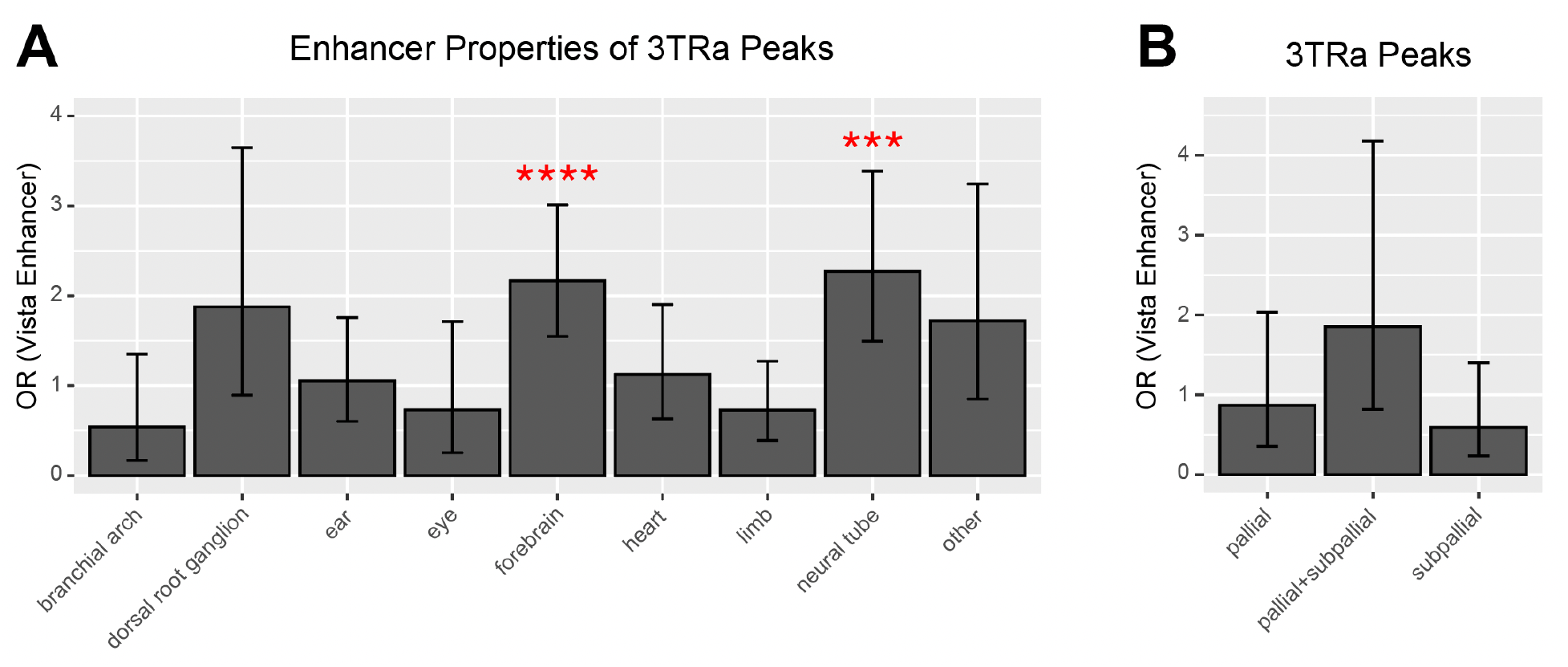
Overlap between ≥3TRa peaks and validated vista enhancers. **A)** The odds of VISTA enhancer activity in a given tissue based on the presence of a distal ≥3TRa peak. ***FDR<0.001, ****FDR<0.0001. **B)** The analysis in ‘A’ is repeated for a subset of VISTA enhancer sequences reclassified by expression patterns in the pallium and subpallium (Table S6).

**Figure S9.**
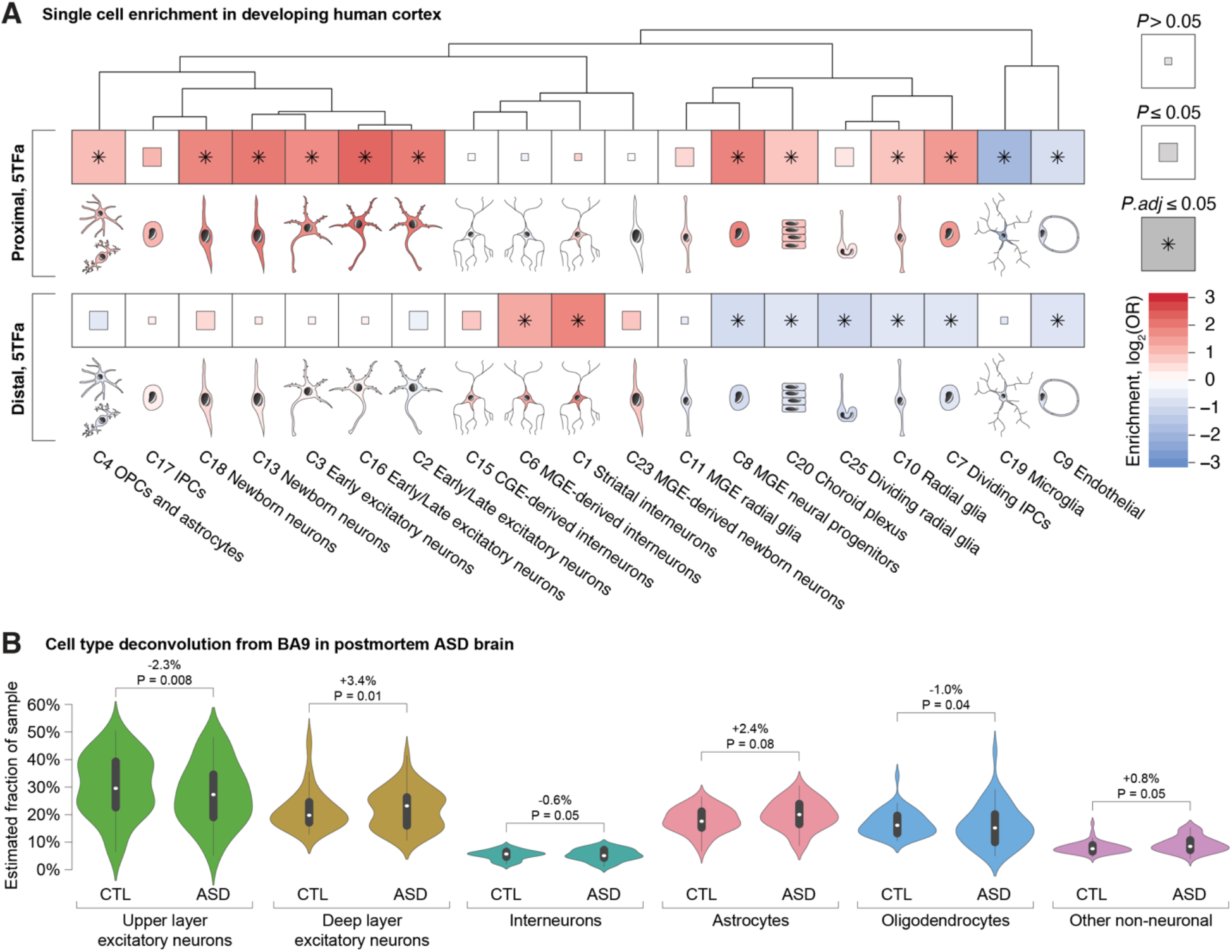
Cell type enrichment and functional consequences. **A)** Cell type clusters from the human fetal cortex (*17*) were assessed for enrichment of genes targeted by all five ASD-associated TRs and ATAC-seq (5TRa) in human (GW23) fetal cortex. The degree of enrichment is indicated by color; significance threshold is indicated by box size/asterisk. **B)** Cell type proportions by major classes of cell types, as estimated by deconvolution of bulk RNA-seq in postmortem brain from ASD cases and controls (*8*).

**Figure S10.**
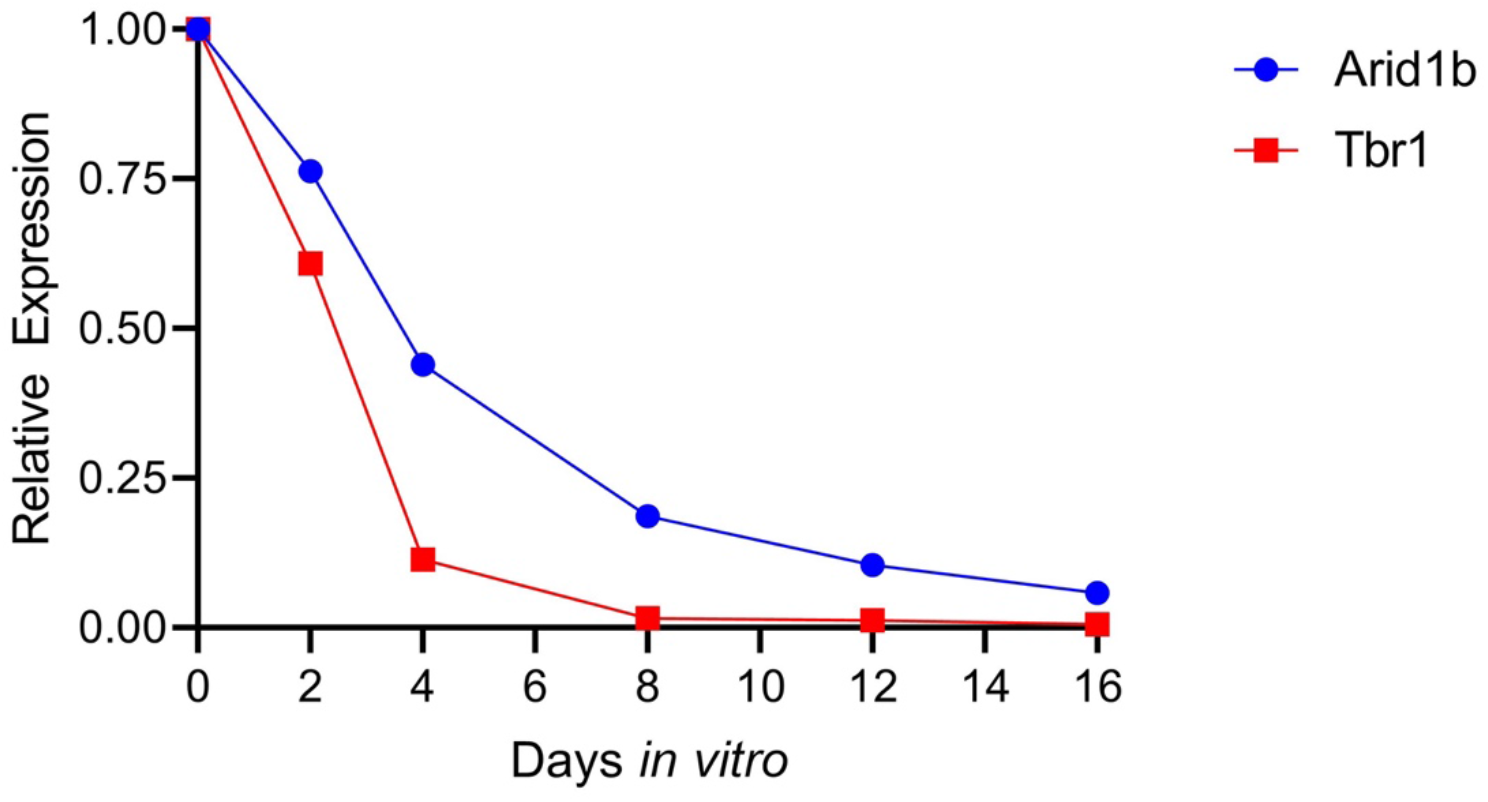
Longitudinal assessment of changes in *Arid1b* and *Tbr1* expression *in vitro* following CRISPRi treatment. CRISPRi guides were designed against promoter regions of *Arid1b* and *Tbr1.* Changes in *Arid1b* and *Tbr1* expression levels were compared to scrambled guides and the *Ef1a* housekeeping gene. CRISPRi reduced relative expression of *Arid1b* to 75% by day 2 and 25% by day 8; likewise, *Tbr1* was reduced to 50% by day 2 and 5% by day 8

## References

1. S. Sandin, P. Lichtenstein, R. Kuja-Halkola, C. Hultman, H. Larsson, A. Reichenberg, The Heritability of Autism Spectrum Disorder. Jama. 318, 1182–1184 (2017).

2. F. K. Satterstrom, J. A. Kosmicki, J. Wang, M. S. Breen, S. De Rubeis, J.-Y. An, M. Peng, R. Collins, J. Grove, L. Klei, C. Stevens, J. Reichert, M. S. Mulhern, M. Artomov, S. Gerges, B. Sheppard, X. Xu, A. Bhaduri, U. Norman, H. Brand, G. Schwartz, R. Nguyen, E. E. Guerrero, C. Dias, B. Aleksic, R. Anney, M. Barbosa, S. Bishop, A. Brusco, J. Bybjerg-Grauholm, A. Carracedo, M. C. Y. Chan, A. G. Chiocchetti, B. H. Y. Chung, H. Coon, M. L. Cuccaro, A. Curró, B. Dalla Bernardina, R. Doan, E. Domenici, S. Dong, C. Fallerini, M. Fernández-Prieto, G. B. Ferrero, C. M. Freitag, M. Fromer, J. J. Gargus, D. Geschwind, E. Giorgio, J. González-Peñas, S. Guter, D. Halpern, E. Hansen-Kiss, X. He, G. E. Herman, I. Hertz-Picciotto, D. M. Hougaard, C. M. Hultman, I. Ionita-Laza, S. Jacob, J. Jamison, A. Jugessur, M. Kaartinen, G. P. Knudsen, A. Kolevzon, I. Kushima, S. L. Lee, T. Lehtimäki, E. T. Lim, C. Lintas, W. I. Lipkin, D. Lopergolo, F. Lopes, Y. Ludena, P. Maciel, P. Magnus, B. Mahjani, N. Maltman, D. S. Manoach, G. Meiri, I. Menashe, J. Miller, N. Minshew, E. M. S. Montenegro, D. Moreira, E. M. Morrow, O. Mors, P. B. Mortensen, M. Mosconi, P. Muglia, B. M. Neale, M. Nordentoft, N. Ozaki, A. Palotie, M. Parellada, M. R. Passos-Bueno, M. Pericak-Vance, A. M. Persico, I. Pessah, K. Puura, A. Reichenberg, A. Renieri, E. Riberi, E. B. Robinson, K. E. Samocha, S. Sandin, S. L. Santangelo, G. Schellenberg, S. W. Scherer, S. Schlitt, R. Schmidt, L. Schmitt, I. M. W. Silva, T. Singh, P. M. Siper, M. Smith, G. Soares, C. Stoltenberg, P. Suren, E. Susser, J. Sweeney, P. Szatmari, L. Tang, F. Tassone, K. Teufel, E. Trabetti, M. D. P. Trelles, C. A. Walsh, L. A. Weiss, T. Werge, D. M. Werling, E. M. Wigdor, E. Wilkinson, A. J. Willsey, T. W. Yu, M. H. C. Yu, R. Yuen, E. Zachi, E. Agerbo, T. D. Als, V. Appadurai, M. Bækvad-Hansen, R. Belliveau, A. Buil, C. E. Carey, F. Cerrato, K. Chambert, C. Churchhouse, S. Dalsgaard, D. Demontis, A. Dumont, J. Goldstein, C. S. Hansen, M. E. Hauberg, M. V. Hollegaard, D. P. Howrigan, H. Huang, J. Maller, A. R. Martin, J. Martin, M. Mattheisen, J. Moran, J. Pallesen, D. S. Palmer, C. B. Pedersen, M. G. Pedersen, T. Poterba, J. B. Poulsen, S. Ripke, A. J. Schork, W. K. Thompson, P. Turley, R. K. Walters, C. Betancur, E. H. Cook, L. Gallagher, M. Gill, J. S. Sutcliffe, A. Thurm, M. E. Zwick, A. D. Børglum, M. W. State, A. E. Cicek, M. E. Talkowski, D. J. Cutler, B. Devlin, S. J. Sanders, K. Roeder, M. J. Daly, J. D. Buxbaum, Large-Scale Exome Sequencing Study Implicates Both Developmental and Functional Changes in the Neurobiology of Autism. Cell. 180 (2020), doi:10.1016/j.cell.2019.12.036.

3. J. M. Fu, F. K. Satterstrom, M. Peng, H. Brand, R. L. Collins, S. Dong, L. Klei, C. R. Stevens, C. Cusick, M. Babadi, E. Banks, B. Collins, S. Dodge, S. B. Gabriel, L. Gauthier, S. K. Lee, L. Liang, A. Ljungdahl, B. Mahjani, L. Sloofman, A. Smirnov, M. Barbosa, A. Brusco, B. H. Y. Chung, M. L. Cuccaro, E. Domenici, G. B. Ferrero, J. J. Gargus, G. E. Herman, I. Hertz-Picciotto, P. Maciel, D. S. Manoach, M. R. Passos-Bueno, A. M. Persico, A. Renieri, F. Tassone, E. Trabetti, G. Campos, M. C. Y. Chan, C. Fallerini, E. Giorgio, A. C. Girard, E. Hansen-Kiss, S. L. Lee, C. Lintas, Y. Ludena, R. Nguyen, L. Pavinato, M. Pericak-Vance, I. Pessah, E. Riberi, R. Schmidt, M. Smith, C. I. C. Souza, S. Trajkova, J. Y. T. Wang, M. H. C. Yu, T. A. S. C. (ASC), B. I. C. for C. D. G. (Broad-CCDG), iPSYCH-B. Consortium, D. J. Cutler, S. De Rubeis, J. D. Buxbaum, M. J. Daly, B. Devlin, K. Roeder, S. J. Sanders, M. E. Talkowski, medRxiv, in press, doi:10.1101/2021.12.20.21267194.

4. A. J. Willsey, S. J. Sanders, M. Li, S. Dong, A. T. Tebbenkamp, R. A. Muhle, S. K. Reilly, L. Lin, S. Fertuzinhos, J. A. Miller, M. T. Murtha, C. Bichsel, W. Niu, J. Cotney, A. G. Ercan-Sencicek, J. Gockley, A. R. Gupta, W. Han, X. He, E. J. Hoffman, L. Klei, J. Lei, W. Liu, L. Liu, C. Lu, X. Xu, Y. Zhu, S. M. Mane, E. S. Lein, L. Wei, J. P. Noonan, K. Roeder, B. Devlin, N. Sestan, M. W. State, Coexpression networks implicate human midfetal deep cortical projection neurons in the pathogenesis of autism. Cell. 155, 997–1007 (2013).

5. N. N. Parikshak, R. Luo, A. Zhang, H. Won, J. K. Lowe, V. Chandran, S. Horvath, D. H. Geschwind, Integrative functional genomic analyses implicate specific molecular pathways and circuits in autism. Cell. 155, 1008–1021 (2013).

6. D. M. Werling, S. Pochareddy, J. Choi, J.-Y. Y. An, B. Sheppard, M. Peng, Z. Li, C. Dastmalchi, G. Santpere, A. M. M. A. M. M. Sousa, A. T. N. N. Tebbenkamp, N. Kaur, F. O. Gulden, M. S. Breen, L. Liang, M. C. Gilson, X. Zhao, S. Dong, L. Klei, A. E. Cicek, J. D. Buxbaum, H. Adle-Biassette, J.-L. L. Thomas, K. A. Aldinger, D. R. O’Day, I. A. Glass, N. A. Zaitlen, M. E. Talkowski, K. Roeder, M. W. State, B. Devlin, S. J. Sanders, N. Sestan, Whole-Genome and RNA Sequencing Reveal Variation and Transcriptomic Coordination in the Developing Human Prefrontal Cortex. Cell Rep. 31, 107489 (2020).

7. N. N. Parikshak, V. Swarup, T. G. Belgard, M. Irimia, G. Ramaswami, M. J. Gandal, C. Hartl, V. Leppa, L. de la T. Ubieta, J. Huang, J. K. Lowe, B. J. Blencowe, S. Horvath, D. H. Geschwind, Genome-wide changes in lncRNA, splicing, and regional gene expression patterns in autism. Nature. 540, 423–427 (2016).

8. J. R. Haney, B. Wamsley, G. T. Chen, S. Parhami, P. S. Emani, N. Chang, G. D. Hoftman, D. de Alba, G. Kale, G. Ramaswami, C. L. Hartl, T. Jin, D. Wang, J. Ou, Y. E. Wu, N. N. Parikshak, V. Swarup, T. G. Belgard, M. Gerstein, B. Pasaniuc, M. J. Gandal, D. H. Geschwind, bioRxiv, in press, doi:10.1101/2020.12.17.423129.

9. D. Velmeshev, L. Schirmer, D. Jung, M. Haeussler, Y. Perez, S. Mayer, A. Bhaduri, N. Goyal, D. H. H. Rowitch, A. R. R. Kriegstein, Single-cell genomics identifies cell type–specific molecular changes in autism. Science (80-.). 364, 685–689 (2019).

10. X. Jin, S. K. Simmons, A. X. Guo, A. S. Shetty, M. Ko, L. Nguyen, E. Robinson, P. Oyler, N. Curry, G. Deangeli, S. Lodato, J. Z. Levin, A. Regev, F. Zhang, P. Arlotta, V. Jokhi, E. Robinson, P. Oyler, N. Curry, G. Deangeli, S. Lodato, J. Z. Levin, A. Regev, F. Zhang, P. Arlotta, V. Jokhi, E. Robinson, P. Oyler, N. Curry, G. Deangeli, S. Lodato, J. Z. Levin, A. Regev, F. Zhang, P. Arlotta, in vivo Perturb-Seq reveals neuronal and glial abnormalities associated with Autism risk genes. Science. 370, 791525 (2020).

11. J. Cotney, R. a Muhle, S. J. Sanders, L. Liu, A. J. Willsey, W. Niu, W. Liu, L. Klei, J. Lei, J. Yin, S. K. Reilly, A. T. Tebbenkamp, C. Bichsel, M. Pletikos, N. Sestan, K. Roeder, M. W. State, B. Devlin, J. P. Noonan, The autism-associated chromatin modifier CHD8 regulates other autism risk genes during human neurodevelopment. Nat Commun. 6, 6404 (2015).

12. A. Sugathan, M. Biagioli, C. Golzio, S. Erdin, I. Blumenthal, P. Manavalan, A. Ragavendran, H. Brand, D. Lucente, J. Miles, S. D. Sheridan, A. Stortchevoi, M. Kellis, S. J. Haggarty, N. Katsanis, J. F. Gusella, M. E. Talkowski, CHD8 regulates neurodevelopmental pathways associated with autism spectrum disorder in neural progenitors. Proc Natl Acad Sci U S A. 111, E4468–77 (2014).

13. J. H. Notwell, W. E. Heavner, S. F. Darbandi, S. Katzman, W. L. Mckenna, C. F. Ortiz-Londono, D. Tastad, M. J. Eckler, J. L. R. Rubenstein, S. K. Mcconnell, B. Chen, G. Bejerano, TBR1 Regulates Autism Risk Genes in the Developing Neocortex. Cold Spring Harb. Lab. Press June. 23, 1–10 (2016).

14. E. Markenscoff-Papadimitriou, F. Binyameen, S. Whalen, J. Price, K. Lim, A. R. Ypsilanti, R. Catta-Preta, E. L.-L. Pai, X. Mu, D. Xu, K. S. Pollard, A. S. Nord, M. W. State, J. L. Rubenstein, Autism risk gene POGZ promotes chromatin accessibility and expression of clustered synaptic genes. Cell Rep. 37, 110089 (2021).

15. F. Aguet, A. A. Brown, S. E. Castel, J. R. Davis, Y. He, B. Jo, P. Mohammadi, Y. S. Park, P. Parsana, A. V. Segrè, B. J. Strober, Z. Zappala, B. B. Cummings, E. T. Gelfand, K. Hadley, K. H. Huang, M. Lek, X. Li, J. L. Nedzel, D. Y. Nguyen, M. S. Noble, T. J. Sullivan, T. Tukiainen, D. G. MacArthur, G. Getz, A. Addington, P. Guan, S. Koester, A. R. Little, N. C. Lockhart, H. M. Moore, A. Rao, J. P. Struewing, S. Volpi, L. E. Brigham, R. Hasz, M. Hunter, C. Johns, M. Johnson, G. Kopen, W. F. Leinweber, J. T. Lonsdale, A. McDonald, B. Mestichelli, K. Myer, B. Roe, M. Salvatore, S. Shad, J. A. Thomas, G. Walters, M. Washington, J. Wheeler, J. Bridge, B. A. Foster, B. M. Gillard, E. Karasik, R. Kumar, M. Miklos, M. T. Moser, S. D. Jewell, R. G. Montroy, D. C. Rohrer, D. Valley, D. C. Mash, D. A. Davis, L. Sobin, M. E. Barcus, P. A. Branton, N. S. Abell, B. Balliu, O. Delaneau, L. Frésard, E. R. Gamazon, D. Garrido-Martín, A. D. H. Gewirtz, G. Gliner, M. J. Gloudemans, B. Han, A. Z. He, F. Hormozdiari, X. Li, B. Liu, E. Y. Kang, I. C. McDowell, H. Ongen, J. J. Palowitch, C. B. Peterson, G. Quon, S. Ripke, A. Saha, A. A. Shabalin, T. C. Shimko, J. H. Sul, N. A. Teran, E. K. Tsang, H. Zhang, Y. H. Zhou, C. D. Bustamante, N. J. Cox, R. Guigó, M. Kellis, M. I. McCarthy, D. F. Conrad, E. Eskin, G. Li, A. B. Nobel, C. Sabatti, B. E. Stranger, X. Wen, F. A. Wright, K. G. Ardlie, E. T. Dermitzakis, T. Lappalainen, A. Battle, C. D. Brown, B. E. Engelhardt, S. B. Montgomery, R. E. Handsaker, S. Kashin, K. J. Karczewski, D. T. Nguyen, C. A. Trowbridge, R. Barshir, O. Basha, G. K. Bogu, L. S. Chen, C. Chiang, F. N. Damani, P. G. Ferreira, I. M. Hall, C. Howald, H. K. Im, Y. Kim, S. Kim-Hellmuth, S. Mangul, J. Monlong, M. Muñoz-Aguirre, A. W. Ndungu, D. L. Nicolae, M. Oliva, N. Panousis, P. Papasaikas, A. J. Payne, J. Quan, F. Reverter, M. Sammeth, A. J. Scott, R. Sodaei, M. Stephens, S. Urbut, M. Van De Bunt, G. Wang, H. S. Xi, E. Yeger-Lotem, J. B. Zaugg, J. M. Akey, D. Bates, J. Chan, M. Claussnitzer, K. Demanelis, M. Diegel, J. A. Doherty, A. P. Feinberg, M. S. Fernando, J. Halow, K. D. Hansen, E. Haugen, P. F. Hickey, L. Hou, F. Jasmine, R. Jian, L. Jiang, A. Johnson, R. Kaul, M. G. Kibriya, K. Lee, J. B. Li, Q. Li, J. Lin, S. Lin, S. Linder, C. Linke, Y. Liu, M. T. Maurano, B. Molinie, J. Nelson, F. J. Neri, Y. Park, B. L. Pierce, N. J. Rinaldi, L. F. Rizzardi, R. Sandstrom, A. Skol, K. S. Smith, M. P. Snyder, J. Stamatoyannopoulos, H. Tang, L. Wang, M. Wang, N. Van Wittenberghe, F. Wu, R. Zhang, C. R. Nierras, L. J. Carithers, J. B. Vaught, S. E. Gould, N. C. Lockart, C. Martin, A. M. Addington, S. E. Koester, A. H. Undale, A. M. Smith, D. E. Tabor, N. V. Roche, J. A. McLean, N. Vatanian, K. L. Robinson, K. M. Valentino, L. Qi, S. Hunter, P. Hariharan, S. Singh, K. S. Um, T. Matose, M. M. Tomaszewski, L. K. Barker, M. Mosavel, L. A. Siminoff, H. M. Traino, P. Flicek, T. Juettemann, M. Ruffier, D. Sheppard, K. Taylor, S. J. Trevanion, D. R. Zerbino, B. Craft, M. Goldman, M. Haeussler, W. J. Kent, C. M. Lee, B. Paten, K. R. Rosenbloom, J. Vivian, J. Zhu, Genetic effects on gene expression across human tissues. Nature. 550, 204– 213 (2017).

16. M. Uhlén, L. Fagerberg, B. M. Hallström, C. Lindskog, P. Oksvold, A. Mardinoglu, Å. Sivertsson, C. Kampf, E. Sjöstedt, A. Asplund, I. Olsson, K. Edlund, E. Lundberg, S. Navani, C. A.-K. Szigyarto, J. Odeberg, D. Djureinovic, J. O. Takanen, S. Hober, T. Alm, P.-H. Edqvist, H. Berling, H. Tegel, J. Mulder, J. Rockberg, P. Nilsson, J. M. Schwenk, M. Hamsten, K. von Feilitzen, M. Forsberg, L. Persson, F. Johansson, M. Zwahlen, G. von Heijne, J. Nielsen, F. Pontén, Proteomics. Tissue-based map of the human proteome. Science. 347, 1260419 (2015).

17. T. J. Nowakowski, A. Bhaduri, A. A. Pollen, B. Alvarado, M. A. Mostajo-Radji, E. Di Lullo, M. Haeussler, C. Sandoval-Espinosa, S. J. Liu, D. Velmeshev, J. R. Ounadjela, J. Shuga, X. Wang, D. A. Lim, J. A. West, A. A. Leyrat, W. J. Kent, A. R. Kriegstein, Spatiotemporal gene expression trajectories reveal developmental hierarchies of the human cortex. Science. 358, 1318–1323 (2017).

18. L. Schirmer, D. Velmeshev, S. Holmqvist, M. Kaufmann, S. Werneburg, D. Jung, S. Vistnes, J. H. Stockley, A. Young, M. Steindel, B. Tung, N. Goyal, A. Bhaduri, S. Mayer, J. B. Engler, O. A. Bayraktar, R. J. M. Franklin, M. Haeussler, R. Reynolds, D. P. Schafer, M. A. Friese, L. R. Shiow, A. R. Kriegstein, D. H. Rowitch, Neuronal vulnerability and multilineage diversity in multiple sclerosis. Nature. 573, 75–82 (2019).

19. M. Li, G. Santpere, Y. I. Kawasawa, O. V. Evgrafov, F. O. Gulden, S. Pochareddy, S. M. Sunkin, Z. Li, Y. Shin, Y. Zhu, A. M. M. Sousa, D. M. Werling, R. R. Kitchen, H. J. Kang, M. Pletikos, J. Choi, S. Muchnik, X. Xu, D. Wang, B. Lorente-Galdos, S. Liu, P. Giusti-Rodríguez, H. Won, C. A. De Leeuw, A. F. Pardiñas, M. A. Reimers, A. J. Willsey, A. Oldre, A. Szafer, A. Camarena, A. Cherskov, A. W. Charney, A. Abyzov, A. Kozlenkov, A. Safi, A. R. Jones, A. Ashley-Koch, A. Ebbert, A. J. Price, A. Sekijima, A. Kefi, A. Bernard, A. Amiri, A. Sboner, A. Clark, A. E. Jaffe, A. T. N. Tebbenkamp, A. J. Sodt, A. Guillozet-Bongaarts, A. C. Nairn, A. Carey, A. Huttner, A. Chervenak, A. Szekely, A. W. Shieh, A. Harmanci, B. K. Lipska, B. C. Carlyle, B. W. Gregor, B. S. Kassim, B. Sheppard, C. Bichsel, C. G. Hahn, C. K. Lee, C. Chen, C. L. Kuan, C. Dang, C. Lau, C. Cuhaciyan, C. Armoskus, C. E. Mason, C. Liu, C. R. Slaughterbeck, C. Bennet, D. Pinto, D. Polioudakis, D. Franjic, D. J. Miller, D. Bertagnolli, D. A. Lewis, D. Feng, D. Sandman, D. Clarke, D. Williams, D. DelValle, D. Fitzgerald, E. H. Shen, E. Flatow, E. Zharovsky, E. E. Burke, E. Olson, E. Fulfs, E. Mattei, E. Hadjimichael, E. Deelman, F. C. P. Navarro, F. Wu, F. Lee, F. Cheng, F. S. Goes, F. M. Vaccarino, F. Liu, G. E. Hoffman, G. Gürsoy, G. Gee, G. Mehta, G. Coppola, G. Giase, G. Sedmak, G. D. Johnson, G. A. Wray, G. E. Crawford, G. Gu, H. van Bakel, H. Witt, H. J. Yoon, H. Pratt, H. Zhao, I. A. Glass, J. Huey, J. Arnold, J. P. Noonan, J. Bendl, J. M. Jochim, J. Goldy, J. Herstein, J. R. Wiseman, J. A. Miller, J. Mariani, J. Stoll, J. Moore, J. Szatkiewicz, J. Leng, J. Zhang, J. Parente, J. Rozowsky, J. F. Fullard, J. G. Hohmann, J. Morris, J. W. Phillips, J. Warrell, J. H. Shin, J. Y. An, J. Belmont, J. Nyhus, J. Pendergraft, J. Bryois, K. Roll, K. S. Grennan, K. Aiona, K. P. White, K. A. Aldinger, K. A. Smith, K. Girdhar, K. Brouner, L. M. Mangravite, L. Brown, L. Collado-Torres, L. Cheng, L. Gourley, L. Song, L. T. Ubieta, L. Habegger, L. Ng, M. E. Hauberg, M. Onorati, M. J. Webster, M. Kundakovic, M. Skarica, M. B. Johnson, M. M. Chen, M. E. Garrett, M. Sarreal, M. Reding, M. Gu, M. A. Peters, M. Fisher, M. J. Gandal, M. Purcaro, M. Smith, M. Brown, M. Shibata, M. Xu, M. Yang, M. Ray, N. V. Shapovalova, N. Francoeur, N. Sjoquist, N. Mastan, N. Kaur, N. Parikshak, N. F. Mosqueda, N. K. Ngo, N. Dee, N. A. Ivanov, O. Devillers, P. Roussos, P. D. Parker, P. Manser, P. Wohnoutka, P. J. Farnham, P. Zandi, P. S. Emani, R. A. Dalley, R. Mayani, R. Tao, R. Gittin, R. E. Straub, R. P. Lifton, R. Jacobov, R. E. Howard, R. B. Park, R. Dai, S. Abramowicz, S. Akbarian, S. Schreiner, S. Ma, S. E. Parry, S. Shapouri, S. Weissman, S. Caldejon, S. Mane, S. L. Ding, S. Scuderi, S. Dracheva, S. Butler, S. N. Lisgo, S. K. Rhie, S. Lindsay, S. Datta, T. Souaiaia, T. Roychowdhury, T. Gomez, T. Naluai-Cecchini, T. Beach, T. Goodman, T. Gao, T. A. Dolbeare, T. Fliss, T. E. Reddy, T. Chen, T. Brunetti, T. A. Lemon, T. Desta, T. Borrman, V. Haroutunian, V. N. Spitsyna, V. Swarup, X. Shi, Y. Jiang, Y. Xia, Y. H. Chen, Y. Wang, Y. Chae, Y. T. Yang, Y. Kim, Z. L. Riley, Z. Krsnik, Z. Deng, Z. Weng, Z. Lin, M. Hu, F. Jin, Y. Li, M. J. Owen, M. C. O’Donovan, J. T. R. Walters, D. Posthuma, P. Levitt, D. R. Weinberger, T. M. Hyde, J. E. Kleinman, D. H. Geschwind, M. J. Hawrylycz, M. W. State, S. J. Sanders, P. F. Sullivan, M. B. Gerstein, E. S. Lein, J. A. Knowles, N. Sestan, Integrative functional genomic analysis of human brain development and neuropsychiatric risks. Science. 362 (2018), doi:10.1126/science.aat7615.

20. S. Fazel Darbandi, S. E. Robinson Schwartz, Q. Qi, R. Catta-Preta, E. L.-L. Pai, J. D. Mandell, A. Everitt, A. Rubin, R. A. Krasnoff, S. Katzman, D. Tastad, A. S. Nord, A. J. Willsey, B. Chen, M. W. State, V. S. Sohal, J. L. R. Rubenstein, Neonatal Tbr1 Dosage Controls Cortical Layer 6 Connectivity. Neuron. 100, 831–845.e7 (2018).

21. O. Chodelkova, J. Masek, V. Korinek, Z. Kozmik, O. Machon, Tcf7L2 is essential for neurogenesis in the developing mouse neocortex. Neural Dev. 13, 8 (2018).

22. M. Lee, J. Yoon, H. Song, B. Lee, D. T. Lam, J. Yoon, K. Baek, H. Clevers, Y. Jeong, Tcf7l2 plays crucial roles in forebrain development through regulation of thalamic and habenular neuron identity and connectivity. Dev. Biol. 424, 62–76 (2017).

23. X. Chen, H. Xu, P. Yuan, F. Fang, M. Huss, V. B. Vega, E. Wong, Y. L. Orlov, W. Zhang, J. Jiang, Y. H. Loh, H. C. Yeo, Z. X. Yeo, V. Narang, K. R. Govindarajan, B. Leong, A. Shahab, Y. Ruan, G. Bourque, W. K. Sung, N. D. Clarke, C. L. Wei, H. H. Ng, Integration of External Signaling Pathways with the Core Transcriptional Network in Embryonic Stem Cells. Cell. 133, 1106–1117 (2008).

24. A. S. Hansen, I. Pustova, C. Cattoglio, R. Tjian, X. Darzacq, CTCF and cohesin regulate chromatin loop stability with distinct dynamics. Elife. 6, e25776 (2017).

25. E. Markenscoff-Papadimitriou, S. Whalen, P. Przytycki, R. Thomas, F. Binyameen, T. J. Nowakowski, A. R. Kriegstein, S. J. Sanders, M. W. State, K. S. Pollard, J. L. Rubenstein, A Chromatin Accessibility Atlas of the Developing Human Telencephalon. Cell. 182, 754–769.e18 (2020).

26. J. Kaplanis, K. E. Samocha, L. Wiel, Z. Zhang, K. J. Arvai, R. Y. Eberhardt, G. Gallone, S. H. Lelieveld, H. C. Martin, J. F. McRae, P. J. Short, R. I. Torene, E. de Boer, P. Danecek, E. J. Gardner, N. Huang, J. Lord, I. Martincorena, R. Pfundt, M. R. F. Reijnders, A. Yeung, H. G. Yntema, L. E. L. M. Vissers, J. Juusola, C. F. Wright, H. G. Brunner, H. V Firth, D. R. FitzPatrick, J. C. Barrett, M. E. Hurles, C. Gilissen, K. Retterer, Evidence for 28 genetic disorders discovered by combining healthcare and research data. Nature. 586, 757–762 (2020).

27. T. Singh, T. Poterba, D. Curtis, H. Akil, M. Al Eissa, J. D. Barchas, N. Bass, T. B. Bigdeli, G. Breen, E. J. Bromet, P. F. Buckley, W. E. Bunney, J. Bybjerg-Grauholm, W. F. Byerley, S. B. Chapman, W. J. Chen, C. Churchhouse, N. Craddock, C. M. Cusick, L. DeLisi, S. Dodge, M. A. Escamilla, S. Eskelinen, A. H. Fanous, S. V Faraone, A. Fiorentino, L. Francioli, S. B. Gabriel, D. Gage, S. A. Gagliano Taliun, A. Ganna, G. Genovese, D. C. Glahn, J. Grove, M.-H. Hall, E. Hämäläinen, H. O. Heyne, M. Holi, D. M. Hougaard, D. P. Howrigan, H. Huang, H.-G. Hwu, R. S. Kahn, H. M. Kang, K. J. Karczewski, G. Kirov, J. A. Knowles, F. S. Lee, D. S. Lehrer, F. Lescai, D. Malaspina, S. R. Marder, S. A. McCarroll, A. M. McIntosh, H. Medeiros, L. Milani, C. P. Morley, D. W. Morris, P. B. Mortensen, R. M. Myers, M. Nordentoft, N. L. O’Brien, A. M. Olivares, D. Ongur, W. H. Ouwehand, D. S. Palmer, T. Paunio, D. Quested, M. H. Rapaport, E. Rees, B. Rollins, F. K. Satterstrom, A. Schatzberg, E. Scolnick, L. J. Scott, S. I. Sharp, P. Sklar, J. W. Smoller, J. L. Sobell, M. Solomonson, E. A. Stahl, C. R. Stevens, J. Suvisaari, G. Tiao, S. J. Watson, N. A. Watts, D. H. Blackwood, A. D. Børglum, B. M. Cohen, A. P. Corvin, T. Esko, N. B. Freimer, S. J. Glatt, C. M. Hultman, A. McQuillin, A. Palotie, C. N. Pato, M. T. Pato, A. E. Pulver, D. St Clair, M. T. Tsuang, M. P. Vawter, J. T. Walters, T. M. Werge, R. A. Ophoff, P. F. Sullivan, M. J. Owen, M. Boehnke, M. C. O’Donovan, B. M. Neale, M. J. Daly, Rare coding variants in ten genes confer substantial risk for schizophrenia. Nature. 604, 509–516 (2022).

28. F. K. Satterstrom, J. A. Kosmicki, J. Wang, M. S. Breen, S. De Rubeis, J.-Y. An, M. Peng, R. Collins, J. Grove, L. Klei, C. Stevens, J. Reichert, M. S. Mulhern, M. Artomov, S. Gerges, B. Sheppard, X. Xu, A. Bhaduri, U. Norman, H. Brand, G. Schwartz, R. Nguyen, E. E. Guerrero, C. Dias, C. Betancur, E. H. Cook, L. Gallagher, M. Gill, J. S. Sutcliffe, A. Thurm, M. E. Zwick, A. D. Borglum, M. W. State, A. E. Cicek, M. E. Talkowski, D. J. Cutler, B. Devlin, S. J. Sanders, K. Roeder, M. J. Daly, J. D. Buxbaum, Large-Scale Exome Sequencing Study Implicates Both Developmental and Functional Changes in the Neurobiology of Autism. Cell (2020), doi:10.1016/j.cell.2019.12.036.

29. C. P. Fulco, J. Nasser, T. R. Jones, G. Munson, D. T. Bergman, V. Subramanian, S. R. Grossman, R. Anyoha, B. R. Doughty, T. A. Patwardhan, T. H. Nguyen, M. Kane, E. M. Perez, N. C. Durand, C. A. Lareau, E. K. Stamenova, E. L. Aiden, E. S. Lander, J. M. Engreitz, Activity-by-contact model of enhancer–promoter regulation from thousands of CRISPR perturbations. Nat. Genet. 51, 1664–1669 (2019).

30. H. Won, L. de la Torre-Ubieta, J. L. Stein, N. N. Parikshak, J. Huang, C. K. Opland, M. J. Gandal, G. J. Sutton, F. Hormozdiari, D. Lu, C. Lee, E. Eskin, I. Voineagu, J. Ernst, D. H. Geschwind, Chromosome conformation elucidates regulatory relationships in developing human brain. Nature. 538, 523–527 (2016).

31. A. Visel, S. Minovitsky, I. Dubchak, L. A. Pennacchio, VISTA Enhancer Browser--a database of tissue-specific human enhancers. Nucleic Acids Res. 35, D88–92 (2007).

32. K. Pattabiraman, O. Golonzhka, S. Lindtner, A. S. Nord, L. Taher, R. Hoch, S. N. Silberberg, D. Zhang, B. Chen, H. Zeng, L. A. Pennacchio, L. Puelles, A. Visel, J. L. R. Rubenstein, Transcriptional regulation of enhancers active in protodomains of the developing cerebral cortex. Neuron. 82, 989–1003 (2014).

33. D. Polioudakis, L. de la Torre-Ubieta, J. Langerman, A. G. Elkins, X. Shi, J. L. Stein, C. K. Vuong, S. Nichterwitz, M. Gevorgian, C. K. Opland, D. Lu, W. Connell, E. K. Ruzzo, J. K. Lowe, T. Hadzic, F. I. Hinz, S. Sabri, W. E. Lowry, M. B. Gerstein, K. Plath, D. H. Geschwind, A Single-Cell Transcriptomic Atlas of Human Neocortical Development during Mid-gestation. Neuron. 103, 785–801.e8 (2019).

34. S. De Rubeis, X. He, A. P. A. A. P. Goldberg, C. S. C. C. S. Poultney, K. Samocha, A. E. Cicek, Y. Kou, L. Liu, M. Fromer, S. Walker, T. T. Singh, L. Klei, J. Kosmicki, F. Shih-Chen, B. Aleksic, M. Biscaldi, P. P. F. Bolton, J. M. J. Brownfeld, J. Cai, N. N. G. Campbell, A. Carracedo, M. H. M. Chahrour, A. A. G. Chiocchetti, H. Coon, E. L. E. E. L. Crawford, S. S. R. Curran, G. Dawson, E. Duketis, B. B. A. Fernandez, L. Gallagher, E. Geller, S. J. S. S. J. Guter, R. S. Hill, J. Ionita-Laza, P. Jimenz Gonzalez, H. Kilpinen, S. M. S. Klauck, A. Kolevzon, I. Lee, I. Lei, J. Lei, T. Lehtimäki, C.-F. F. C. Lin, A. Ma’ayan, C. R. C. Marshall, A. L. McInnes, B. Neale, M. J. M. Owen, N. N. Ozaki, M. Parellada, J. R. J. Parr, S. Purcell, K. Puura, D. Rajagopalan, K. Rehnström, A. Reichenberg, A. Sabo, M. Sachse, S. J. S. S. J. Sanders, C. Schafer, M. Schulte-Rüther, D. Skuse, C. Stevens, P. Szatmari, K. Tammimies, O. Valladares, A. Voran, W. Li-San, L. A. Weiss, A. J. Willsey, T. T. W. Yu, R. R. K. C. Yuen, E. H. E. E. H. Cook, C. M. C. C. M. Freitag, M. Gill, C. M. C. C. M. Hultman, T. Lehner, A. A. Palotie, G. G. D. Schellenberg, P. Sklar, M. M. W. State, J. S. J. Sutcliffe, C. A. C. A. C. Walsh, S. W. S. S. W. Scherer, M. M. E. Zwick, J. C. J. C. Barett, D. J. D. Cutler, K. Roeder, B. Devlin, M. M. J. Daly, J. D. J. Buxbaum, D. D. D. Study, H. M. C. for Autism, U. Consortium, A. Ercument Cicek, Y. Kou, L. Liu, M. Fromer, S. Walker, T. T. Singh, L. Klei, J. Kosmicki, S.-C. Fu, B. Aleksic, M. Biscaldi, P. P. F. Bolton, J. M. J. Brownfeld, J. Cai, N. N. G. Campbell, A. Carracedo, M. H. M. Chahrour, A. A. G. Chiocchetti, H. Coon, E. L. E. E. L. Crawford, L. Crooks, S. S. R. Curran, G. Dawson, E. Duketis, B. B. A. Fernandez, L. Gallagher, E. Geller, S. J. S. S. J. Guter, R. Sean Hill, I. Ionita-Laza, P. Jimenez Gonzalez, H. Kilpinen, S. M. S. Klauck, A. Kolevzon, I. Lee, J. Lei, T. Lehtimäki, C.-F. F. C. Lin, A. Ma’ayan, C. R. C. Marshall, A. L. McInnes, B. Neale, M. J. M. Owen, N. N. Ozaki, M. Parellada, J. R. J. Parr, S. Purcell, K. Puura, D. Rajagopalan, K. Rehnström, A. Reichenberg, A. Sabo, M. Sachse, S. J. S. S. J. Sanders, C. Schafer, M. Schulte-Rüther, D. Skuse, C. Stevens, P. Szatmari, K. Tammimies, O. Valladares, A. Voran, L.-S. Wang, L. A. Weiss, A. Jeremy Willsey, T. T. W. Yu, R. R. K. C. Yuen, E. H. E. E. H. Cook, C. M. C. C. M. Freitag, M. Gill, C. M. C. C. M. Hultman, T. Lehner, A. A. Palotie, G. G. D. Schellenberg, P. Sklar, M. M. W. State, J. S. J. Sutcliffe, C. A. C. A. C. Walsh, S. W. S. S. W. Scherer, M. M. E. Zwick, J. C. Barrett, D. J. D. Cutler, K. Roeder, B. Devlin, M. M. J. Daly, J. D. J. Buxbaum, P. Jimenz, Gonzalez, H. Kilpinen, S. M. S. Klauck, A. Kolevzon, I. Lee, I. Lei, J. Lei, T. Lehtimäki, C.-F. F. C. Lin, A. Ma’ayan, C. R. C. Marshall, A. L. McInnes, B. Neale, M. J. M. Owen, N. N. Ozaki, M. Parellada, J. R. J. Parr, S. Purcell, K. Puura, D. Rajagopalan, K. Rehnström, A. Reichenberg, A. Sabo, M. Sachse, S. J. S. S. J. Sanders, C. Schafer, M. Schulte-Rüther, D. Skuse, C. Stevens, P. Szatmari, K. Tammimies, O. Valladares, A. Voran, W. Li-San, L. A. Weiss, A. J. Willsey, T. T. W. Yu, R. R. K. C. Yuen, D. D. D. Study, H. M. C. for Autism, U. Consortium, E. H. E. E. H. Cook, C. M. C. C. M. Freitag, M. Gill, C. M. C. C. M. Hultman, T. Lehner, A. A. Palotie, G. G. D. Schellenberg, P. Sklar, M. M. W. State, J. S. J. Sutcliffe, C. A. C. A. C. Walsh, S. W. S. S. W. Scherer, M. M. E. Zwick, J. C. J. C. Barett, D. J. D. Cutler, K. Roeder, B. Devlin, M. M. J. Daly, J. D. J. Buxbaum, DDD Study, Homozygosity Mapping Collaborative for Autism, UK10K Consortium, E. H. E. E. H. Cook, C. M. C. C. M. Freitag, M. Gill, C. M. C. C. M. Hultman, T. Lehner, A. A. Palotie, G. G. D. Schellenberg, P. Sklar, M. M. W. State, J. S. J. Sutcliffe, C. A. C. A. C. Walsh, S. W. S. S. W. Scherer, M. M. E. Zwick, J. C. J. C. Barett, D. J. D. Cutler, K. Roeder, B. Devlin, M. M. J. Daly, J. D. J. Buxbaum, Synaptic, transcriptional and chromatin genes disrupted in autism. Nature. 515, 209–215 (2014).

35. S. J. Sanders, X. He, A. J. Willsey, B. Devlin, K. Roeder, M. W. State, S. J. Sanders, X. He, A. J. Willsey, A. G. Ercan-sencicek, K. E. Samocha, A. E. Cicek, M. T. Murtha, V. H. Bal, S. L. Bishop, S. Dong, A. P. Goldberg, C. Jinlu, J. F. Keaney, L. Klei, J. D. Mandell, D. Moreno-De-Luca, C. S. Poultney, E. B. Robinson, L. Smith, T. Solli-Nowlan, M. Y. Su, N. A. Teran, M. F. Walker, D. M. Werling, A. L. Beaudet, R. M. Cantor, E. Fombonne, D. H. Geschwind, D. E. Grice, C. Lord, J. K. Lowe, S. M. Mane, D. M. Martin, E. M. Morrow, M. E. Talkowski, J. S. Sutcliffe, C. A. Walsh, T. W. Yu, D. H. Ledbetter, C. L. Martin, E. H. Cook, J. D. Buxbaum, M. J. Daly, B. Devlin, K. Roeder, M. W. State, A. S. Consortium, Autism Sequencing Consortium, D. H. Ledbetter, C. L. Martin, E. H. Cook, J. D. Buxbaum, M. J. Daly, B. Devlin, K. Roeder, M. W. State, Insights into Autism Spectrum Disorder Genomic Architecture and Biology from 71 Risk Loci. Neuron. 87, 1215–1233 (2015).

36. A. E. Trevino, F. Müller, J. Andersen, L. Sundaram, A. Kathiria, A. Shcherbina, K. Farh, H. Y. Chang, A. M. Pașca, A. Kundaje, S. P. Pașca, W. J. Greenleaf, Chromatin and gene-regulatory dynamics of the developing human cerebral cortex at single-cell resolution. Cell. 184, 5053–5069.e23 (2021).

37. J. den Hoed, E. Sollis, H. Venselaar, S. B. Estruch, P. Deriziotis, S. E. Fisher, Functional characterization of TBR1 variants in neurodevelopmental disorder. Sci. Rep. 8, 14279 (2018).

38. P. Deriziotis, B. J. O’Roak, S. A. Graham, S. B. Estruch, D. Dimitropoulou, R. A. Bernier, J. Gerdts, J. Shendure, E. E. Eichler, S. E. Fisher, De novo TBR1 mutations in sporadic autism disrupt protein functions. Nat. Commun. 5, 4954 (2014).

39. S. B. Estruch, S. A. Graham, M. Quevedo, A. Vino, D. H. W. Dekkers, P. Deriziotis, E. Sollis, J. Demmers, R. A. Poot, S. E. Fisher, Proteomic analysis of FOXP proteins reveals interactions between cortical transcription factors associated with neurodevelopmental disorders. Hum. Mol. Genet. 27, 1212–1227 (2018).

40. M. P. Walker, C. M. Stopford, M. Cederlund, F. Fang, C. Jahn, A. D. Rabinowitz, D. Goldfarb, D. M. Graham, F. Yan, A. M. Deal, Y. Fedoriw, K. L. Richards, I. J. Davis, G. Weidinger, B. Damania, M. B. Major, FOXP1 potentiates Wnt/β-catenin signaling in diffuse large B cell lymphoma. Sci. Signal. 8, ra12 (2015).

41. M. Sen, X. Wang, F. H. Hamdan, J. Rapp, J. Eggert, R. L. Kosinsky, F. Wegwitz, A. P. Kutschat, F. S. Younesi, J. Gaedcke, M. Grade, E. Hessmann, A. Papantonis, P. Strӧbel, S. A. Johnsen, ARID1A facilitates KRAS signaling-regulated enhancer activity in an AP1-dependent manner in colorectal cancer cells. Clin. Epigenetics. 11, 92 (2019).

42. R. R. Moody, M.-C. Lo, J. L. Meagher, C.-C. Lin, N. O. Stevers, S. L. Tinsley, I. Jung, A. Matvekas, J. A. Stuckey, D. Sun, Probing the interaction between the histone methyltransferase/deacetylase subunit RBBP4/7 and the transcription factor BCL11A in epigenetic complexes. J. Biol. Chem. 293, 2125–2136 (2018).

43. A. L. Chokas, C. M. Trivedi, M. M. Lu, P. W. Tucker, S. Li, J. A. Epstein, E. E. Morrisey, Foxp1/2/4-NuRD interactions regulate gene expression and epithelial injury response in the lung via regulation of interleukin. J. Biol. Chem. 285, 13304–13313 (2010).

44. J.-Y. An, K. Lin, L. Zhu, D. M. Werling, S. Dong, H. Brand, H. Z. Wang, X. Zhao, G. B. Schwartz, R. L. Collins, B. B. Currall, C. Dastmalchi, J. Dea, C. Duhn, M. C. Gilson, L. Klei, L. Liang, E. Markenscoff-Papadimitriou, S. Pochareddy, N. Ahituv, J. D. Buxbaum, H. Coon, M. J. Daly, Y. S. Kim, G. T. Marth, B. M. Neale, A. R. Quinlan, J. L. Rubenstein, N. Sestan, M. W. State, A. J. Willsey, M. E. Talkowski, B. Devlin, K. Roeder, S. J. Sanders, Genome-wide de novo risk score implicates promoter variation in autism spectrum disorder. Science. 362, eaat6576 (2018).

45. J. Grove, S. Ripke, T. D. Als, M. Mattheisen, R. K. Walters, H. Won, J. Pallesen, E. Agerbo, O. A. Andreassen, R. Anney, S. Awashti, R. Belliveau, F. Bettella, J. D. Buxbaum, J. Bybjerg-Grauholm, M. Bækvad-Hansen, F. Cerrato, K. Chambert, J. H. Christensen, C. Churchhouse, K. Dellenvall, D. Demontis, S. De Rubeis, B. Devlin, S. Djurovic, A. L. Dumont, J. I. Goldstein, C. S. Hansen, M. E. Hauberg, M. V. Hollegaard, S. Hope, D. P. Howrigan, H. Huang, C. M. Hultman, L. Klei, J. Maller, J. Martin, A. R. Martin, J. L. Moran, M. Nyegaard, T. Nærland, D. S. Palmer, A. Palotie, C. B. Pedersen, M. G. Pedersen, T. DPoterba, J. B. Poulsen, B. S. Pourcain, P. Qvist, K. Rehnström, A. Reichenberg, J. Reichert, E. B. Robinson, K. Roeder, P. Roussos, E. Saemundsen, S. Sandin, F. K. Satterstrom, G. Davey Smith, H. Stefansson, S. Steinberg, C. R. Stevens, P. F. Sullivan, P. Turley, G. B. Walters, X. Xu, K. Stefansson, D. H. Geschwind, M. Nordentoft, D. M. Hougaard, T. Werge, O. Mors, P. B. Mortensen, B. M. Neale, M. J. Daly, A. D. Børglum, Identification of common genetic risk variants for autism spectrum disorder. Nat. Genet., 1 (2019).

46. T. Gaugler, L. Klei, S. J. Sanders, C. A. Bodea, A. P. Goldberg, A. B. Lee, M. Mahajan, D. Manaa, Y. Pawitan, J. Reichert, S. Ripke, S. Sandin, P. Sklar, O. Svantesson, A. Reichenberg, C. M. Hultman, B. Devlin, K. Roeder, J. D. Buxbaum, Most genetic risk for autism resides with common variation. Nat Genet. 46, 881–885 (2014).

47. D. J. Araujo, A. G. Anderson, S. Berto, W. Runnels, M. Harper, S. Ammanuel, M. A. Rieger, H. C. Huang, K. Rajkovich, K. W. Loerwald, J. D. Dekker, H. O. Tucker, J. D. Dougherty, J. R. Gibson, G. Konopka, FoxP1 orchestration of ASD-relevant signaling pathways in the striatum. Genes Dev. 29, 2081–2096 (2015).

48. M. J. Gandal, J. R. Haney, N. N. Parikshak, V. Leppa, G. Ramaswami, C. Hartl, A. J. Schork, V. Appadurai, A. Buil, T. M. Werge, C. Liu, K. P. White, S. Horvath, D. H. Geschwind, Shared molecular neuropathology across major psychiatric disorders parallels polygenic overlap. Science. 359, 693–697 (2018).

49. J. S. Fleck, S. M. J. Jansen, D. Wollny, M. Seimiya, F. Zenk, M. Santel, Z. He, J. Gray Camp, B. Treutlein, bioRxiv, in press, doi:10.1101/2021.08.24.457460.

50. F. Bedogni, R. D. Hodge, G. E. Elsen, B. R. Nelson, R. A. M. Daza, R. P. Beyer, T. K. Bammler, J. L. R. Rubenstein, R. F. Hevner, Tbr1 regulates regional and laminar identity of postmitotic neurons in developing neocortex. Proc. Natl. Acad. Sci. U. S. A. 107, 13129– 13134 (2010).

51. S. Fazel Darbandi, S. E. Robinson Schwartz, E. L.-L. Pai, A. Everitt, M. L. Turner, B. N. R. Cheyette, A. J. Willsey, M. W. State, V. S. Sohal, J. L. R. Rubenstein, Enhancing WNT Signaling Restores Cortical Neuronal Spine Maturation and Synaptogenesis in Tbr1 Mutants. Cell Rep. 31, 107495 (2020).

52. B. Tasic, S. Hippenmeyer, C. Wang, M. Gamboa, H. Zong, Y. Chen-Tsai, L. Luo, Proc. Natl. Acad. Sci., in press, doi:10.1073/pnas.1019507108.

53. W. T. Khaled, S. Choon Lee, J. Stingl, X. Chen, H. Raza Ali, O. M. Rueda, F. Hadi, J. Wang, Y. Yu, S.-F. Chin, M. Stratton, A. Futreal, N. A. Jenkins, S. Aparicio, N. G. Copeland, C. J. Watson, C. Caldas, P. Liu, BCL11A is a triple-negative breast cancer gene with critical functions in stem and progenitor cells. Nat. Commun. 6, 5987 (2015).

54. C. Celen, J.-C. Chuang, X. Luo, N. Nijem, A. K. Walker, F. Chen, S. Zhang, A. S. Chung, L. H. Nguyen, I. Nassour, A. Budhipramono, X. Sun, L. A. Bok, M. McEntagart, E. F. Gevers, S. G. Birnbaum, A. J. Eisch, C. M. Powell, W.-P. Ge, G. W. Santen, M. Chahrour, H. Zhu, Arid1b haploinsufficient mice reveal neuropsychiatric phenotypes and reversible causes of growth impairment. Elife. 6 (2017), doi:10.7554/eLife.25730.

55. A. R. Quinlan, I. M. Hall, BEDTools: a flexible suite of utilities for comparing genomic features. Bioinformatics. 26, 841–842 (2010).

56. A. Bulfone, F. Wang, R. Hevner, S. Anderson, T. Cutforth, S. Chen, J. Meneses, R. Pedersen, R. Axel, J. L. Rubenstein, An olfactory sensory map develops in the absence of normal projection neurons or GABAergic interneurons. Neuron. 21, 1273–1282 (1998).

57. R. F. Hevner, L. Shi, N. Justice, Y. Hsueh, M. Sheng, S. Smiga, A. Bulfone, A. M. Goffinet, A. T. Campagnoni, J. L. Rubenstein, Tbr1 regulates differentiation of the preplate and layer 6. Neuron. 29, 353–366 (2001).

58. J.-P. Concordet, M. Haeussler, CRISPOR: intuitive guide selection for CRISPR/Cas9 genome editing experiments and screens. Nucleic Acids Res. 46, W242–W245 (2018).

59. D. Vogt, P.-R. Wu, S. F. Sorrells, C. Arnold, A. Alvarez-Buylla, J. L. R. Rubenstein, Viral-mediated Labeling and Transplantation of Medial Ganglionic Eminence (MGE) Cells for In Vivo Studies. J. Vis. Exp. (2015), doi:10.3791/52740.

60. S. Darbandi, J. P. C. Franck, A comparative study of ryanodine receptor (RyR) gene expression levels in a basal ray-finned fish, bichir (Polypterus ornatipinnis) and the derived euteleost zebrafish (Danio rerio). Comp. Biochem. Physiol. B. Biochem. Mol. Biol. 154, 443–448 (2009).

61. M. W. Pfaffl, A new mathematical model for relative quantification in real-time RT-PCR. Nucleic Acids Res. 29, e45 (2001).

62. A. Visel, L. Taher, H. Girgis, D. May, O. Golonzhka, R. V. Hoch, G. L. McKinsey, K. Pattabiraman, S. N. Silberberg, M. J. Blow, D. V. Hansen, A. S. Nord, J. A. Akiyama, A. Holt, R. Hosseini, S. Phouanenavong, I. Plajzer-Frick, M. Shoukry, V. Afzal, T. Kaplan, A. R. Kriegstein, E. M. Rubin, I. Ovcharenko, L. A. Pennacchio, J. L. R. Rubenstein, A high-resolution enhancer atlas of the developing telencephalon. Cell. 152, 895–908 (2013).

63. C. Soneson, M. I. Love, M. D. Robinson, Differential analyses for RNA-seq: transcript-level estimates improve gene-level inferences. F1000Research. 4, 1521 (2015).

64. M. I. Love, W. Huber, S. Anders, Moderated estimation of fold change and dispersion for RNA-seq data with DESeq2. Genome Biol. 15, 550 (2014).

